# The ontology of the anatomy and development of the solitary ascidian *Ciona*

**DOI:** 10.1101/2020.06.09.140640

**Authors:** Kohji Hotta, Delphine Dauga, Lucia Manni

## Abstract

*Ciona robusta* (*Ciona intestinalis* type A), a model organism for biological studies, belongs to ascidians, the main class of tunicates, which are the closest relatives of vertebrates. In *Ciona*, a project on the ontology of both development and anatomy is ongoing for several years. Its goal is to standardize a resource relating each anatomical structure to developmental stages. Today, the ontology is codified until the hatching larva stage. Here, we present its extension throughout the swimming larva stages, the metamorphosis, until the juvenile stages. For standardizing the developmental ontology, we acquired different time-lapse movies, confocal microscope images and histological serial section images for each developmental event from the hatching larva stage (17.5 hour post fertilization) to the juvenile stage (7 days post fertilization). Combining these data, we defined 12 new distinct developmental stages (from Stage 26 to Stage 37), in addition to the previously defined 26 stages, referred to embryonic development. The new stages were grouped into four Periods named: Adhesion, Tail Absorption, Body Axis Rotation, and Juvenile. To build the anatomical ontology, 203 anatomical entities were identified, defined according to the literature, and annotated, taking advantage from the high resolution and the complementary information obtained from confocal microscopy and histology. The ontology describes the anatomical entities in hierarchical levels, from the cell level (cell lineage) to the tissue/organ level. Comparing the number of entities during development, we found two rounds on entity increase: in addition to the one occurring after fertilization, there is a second one during the Body Axis Rotation Period, when juvenile structures appear. Vice versa, one-third of anatomical entities associated with the embryo/larval life were significantly reduced at the beginning of metamorphosis. Data was finally integrated within the web-based resource “TunicAnatO”, which includes a number of anatomical images and a dictionary with synonyms. This ontology will allow the standardization of data underpinning an accurate annotation of gene expression and the comprehension of mechanisms of differentiation. It will help in understanding the emergence of elaborated structures during both embryogenesis and metamorphosis, shedding light on tissue degeneration and differentiation occurring at metamorphosis.

## Background

Biological data including both spatial and temporal dimensions are essential for understanding the morphological organization of complex structures, such as tissues, organs, and organisms as a whole. Such structures, here called anatomical entities. In the anatomical entity, it constitutes the structural organization of an individual member of a biological species. At cellular level, “the structural organization” is easy to determine, as the organization is limited by the cell membrane. At structural level, an entity could be defined thanks to anatomical particularities (e.g. tail, trunk, atrial cavity) or to a specific function (e.g. heart, brain). Once anatomical entity recognized and defined in a hierarchical way (*i.e.*, organized in an Anatomical Ontology, AO) and put in relationship with a developmental time-table specifying the developmental stage features (*i.e.*, a Developmental Ontology, DO), constitute the basis upon which to build an Anatomical and Developmental Ontology (ADO). The latter is a powerful instrument to standardize different kinds of biological data and an irreplaceable tool associated with model species ^1–5^.

Among tunicates, the sister group of vertebrates ^6,7^, the solitary ascidians *Ciona intestinalis*, are recognized model species for evolutionary, developmental, and ecological studies^8–11^. Recently, it was shown that there were two cryptic species under the name *C.intestinalis*, called types A and B ^12–20^. A taxonomic study ^21^ proposed to rename *C.intestinalis* type A as *Ciona robusta*, and *C.intestinalis* type B as *C.intestinalis* ^21–23^. From an anatomical point of view, very few differences in adults ^22^ and in larvae ^22^ were reported between the two types. In addition, *C. intestinalis* type A (now *C. robusta*) and type B (now *C. intestinalis*) were used as indistinguishable models until 2015. Therefore, all anatomical entities used in this study are common to *Ciona robusta* (*C. intestinalis* type A) and *Ciona intestinalis* (type B), so here the term *Ciona* refers to both species.

In the ascidian larva, the typical chordate body plan can easily be recognized and studied: muscles for tail deflection during swimming flank a notochord; a hollow nerve cord is dorsal to the notochord, whereas an endodermal strand is ventral to it. This makes ascidians a privileged model for understanding the evolution of more complex vertebrates.

For *Ciona*, the ADO so far available regards 26 early developmental stages, from the unfertilized egg (Stage 0) to the hatching larva (Stage 26) ^24^. This ontology is registered in the Bioportal web portal ^25^.

Moreover, representative 3D morphological reconstructions and optic cross-section images implement the ontology and are available in the web-based database FABA (https://www.bpni.bio.keio.ac.jp/chordate/faba/1.4/top.html). In FABA, information about cell lineages in early development was annotated based on previous investigations ^26–29^. Considering that the ascidian embryogenesis is stereotyped, this ontology provides a standardized resource of spatial and temporal information for both *C. robusta* and *C. intestinalis*, as well as other solitary ascidians.

After larval hatching, ascidian larvae disperse, swimming freely and searching for a suitable substrate on which to metamorphose. The metamorphosis is deep and transforms the larva with the chordate body plan into a sessile, filter-feeding adult (Supplementary Figure S1) ^30^. In the latter, the chordate body plan is no longer recognizable, even if some other chordate features, such as the pharyngeal fissures (the stigmata) and the endostyle in the ventral pharynx (homologous to the vertebrate thyroid gland) ^31^, are now visible.

To cover these further developmental phases, we decided to extend the ADO to the post-hatching larva development and metamorphosis. To build up this new part of the ontology, we conducted an anatomical investigation based on complementary methods. The method of phalloidin-staining, successfully used for visualizing anatomical structures until the hatching larva stage ^24^, unfortunately was revealed as less useful, as cells shrink as development proceeds, thereby becoming hardly recognizable. Moreover, in differentiated individuals, low actin-based structures, such as the tunic or pigment cells (otolith and ocellus), are difficult to recognize. Consequently, we produced a comprehensive collection of both confocal scanning laser microscopy (CLSM) of whole-mount specimens, and light microscope images of 1-μm-thick histological serial sections of whole samples, for each developmental stage. For histology, specimens were cut according to the classical planes: transverse, sagittal, and frontal. This allowed us to build a complete anatomical atlas and was necessary for collecting the morphological information related to internal organs as well as the body shape and external surface. Lastly, for each anatomical entity, we annotated its definition, carefully checking the literature since 1893 and considering, in particular, some milestones of ascidian literature, such as the exhaustive description of *C. intestinalis* published by Millar in 1953 ^32^. Because the same anatomical structure was sometimes called with different names by researchers in different periods or belonging to different biological fields, we also annotated synonyms.

All this information, together with stereomicroscopy time-lapse movies, are consultable in the web-based resource called TunicAnatO (<u>Tunic</u>ate <u>Anat</u>omical and developmental <u>O</u>ntology) (https://www.bpni.bio.keio.ac.jp/tunicanato/3.0/). TunicAnatO includes the former FABA database ^24^, therefore covering, in total, 37 developmental stages of *Ciona* development, from the unfertilized egg to the juvenile. Features exhibited in the newly defined Stages 26 to 37 are described in Supplementary Data S2. TunicAnatO is also reachable via the Bioportal (http://bioportal.bioontology.org/), which is the most comprehensive repository of biomedical ontologies, and via the Tunicate Web Portal (http://www.tunicate-portal.org/), which is the main web tool for the Tunicate Community.

## Results

### The DO and AO from the Post-Hatching Larva Stage to the Juvenile Stage: working method

To construct the DO referring to the developmental stages of *Ciona* following the hatching larva stage, time-lapse imaging and CLSM imaging of sequentially fixed specimens of *Ciona robusta* (*Ciona intestinalis* type A) were performed (Fig. 1, Supplementary Files S3-S5). The DO presents the developmental stages grouped in Periods, which in turn are grouped into Meta-Periods, following the conventional nomenclature of ontologies (Table 1). Table 1 shows the 12 newly defined developmental stages, from Stage 26 to Stage 37. Moreover, it also introduces Stage 38-41, here not described. They complete the Juvenile Period of the Post-Metamorphosis Meta-Period (Supplementary Table S3). The 12 new distinct stages correspond to six stages previously described by Chiba and collaborators ^33^. Representative images of individuals belonging to each stage, at both stereomicroscopy and CLSM, were chosen as reference (Fig. 1; Supplementary Video S4 and Supplementary Figure S5).

**Table 1.** *Ciona* developmental stages from the Larva Period to the Juvenile Period. Stages 1-41 were defined in the paper, Stages 1-37 described.

| Stage | Stage name | Characteristics | Time after fertilization<br>18°C<br>(hour post fertilization) | Time after hatch<br>20°C (*2)<br>(hour post hatch) |
| --- | --- | --- | --- | --- |

**Embryonic development, pre-metamorphosis** CirobuD:0000003
| VI. Larva Period (St.26-29, 17.5-24hpf) (*1) |  |  |  |  |  |
| --- | --- | --- | --- | --- | --- |
| St. 26<br>CirobuD:0000049 | hatching larva | Hatching, spherical head shape, immature papillae with pyramidal shape, irregular tail movements | 17hr 30min (17.5hpf) | 0 | St |
| St. 27<br>CirobuD:0000050 | early swimming larva | Spindle-like head shape, regular tail movements and swimming behaviour | 17.5-20 hpf | - | St |
| St. 28<br>CirobuD:0000051 | mid swimming larva | Elongated papillae and expansion of their basal part, squared head, spherical test cells, cilia in epidermal sensory neurons recognizable, preoral lobe recognizable | 20-22hpf | - | St |
| St. 29<br>CirobuD:0000052 | late swimming larva | Longer and narrower head with respect to St. 28, trunk profile squared at transition between trunk and tail | 22-24hpf | - | St |

**Metamorphosis** CirobuD:0000004
| VII. Adhesion period (St.30-31, 24-27hpf) |  |  |  |  |  |
| --- | --- | --- | --- | --- | --- |
| St. 30<br>CirobuD:0000053 | adhesion | Curved papillae, otolith and ocellus remnants recognizable | 24-27 hpf | 6.5-9.5hph | : |
| VIII. Tail absorption period (St.32-34, 27-30hpf) |  |  |  |  |  |
| St. 31<br>CirobuD:0000054 | early tail absorption | Beginning of tail absorption, tail bending at the transition between trunk and tail, otolith and ocellus remnants recognizable. | 27 hpf | avg. 9.9hph | St |
| St. 32<br>CirobuD:0000055 | mid tail absorption | 50% of tail absorbed into trunk; tail shrinked and thickened, otolith and ocellus remnants recognizable | 28 hpf | avg. 10.4hph | St |
| St. 33<br>CirobuD:0000056 | late tail absorption | Tail completely resorbed, papillae no more recognizable, otolith and ocellus remnants recognizable | 29 hpf | avg. 11.4hph | St |
| IX. Body axis rotation period (St.35-37, 30-60hpf) |  |  |  |  |  |
| St. 34<br>CirobuD:0000057 | early body axis rotation | Beginning of body axis rotation (angle between the stalk and the endostyle more than 0°), outer tunic compartment and outer cuticle layer no more present, tunic cells recognizable in definitive tunic, otolith and ocellus remnants recognizable. | 30-36 hpf | avg. 17.7hph | St |
| St. 35<br>CirobuD:0000058 | mid body axis rotation | Body axis rotation of about 30°-60°, one pair of gill-slit recognizable, otolith and ocellus remnants recognizable | 36-45 hpf | avg. 27hph | St |
| St. 36<br>CirobuD:0000059 | late body axis rotation | Two pairs of gill-slit open, body axis rotation of about 80°-90°, filtering and feeding activity present, otolith and ocellus remnants recognizable, heart beating | 45-60 hpf | avg. 37.5hph | St |

**Post-metamorphosis** CirobuD:0000005
| X. Juvenile period (St.37-41, 60hpf~) |  |  |  |  |  |
| --- | --- | --- | --- | --- | --- |
| St. 37<br>CirobuD:0000060 | early juvenile I | Body axis rotation completed, stomach swollen, otolith and ocellus remnants recognizable | 63-72 hpf (3dpf) | 45.5-54.5hph | St |
| St. 38<br>CirobuD:0000061 | early juvenile II | Larval tail remnants totally adsorbed | 3-4 dpf | 3dph | St |
| St. 39<br>CirobuD:0000062 | mid juvenile I | Additional gill slit begin to open, appearance of stomach, gut and neural grand | 4-6 dpf | 5dph | St |
| St.40<br>CirobuD:0000063 | mid juvenile II | Gonad in form of oval vesicle (corresponding to Stage 6 in Chiba et. al., 2004) | 6-7dpf | 6dph | St |
| St.41<br>CirobuD:0000064 | late juvenile | Atrial siphon begins to fuse (corresponding to Stage 7 in Chiba et. al., 2004) | 7dpf~ | 6dph~ | St |

\*1 The duration of larval swimming differs among individuals. Matsunobu et al. (2015) showed that the hatched larva requires at least three or four hours to get competence to commence metamorphosis. The time after fertilization during Larva Period was broad.
\*2 The average timecourse of Tail absorption period and Body axis rotation period in same batch was set-up once again after hatching.

**Figure 1.**
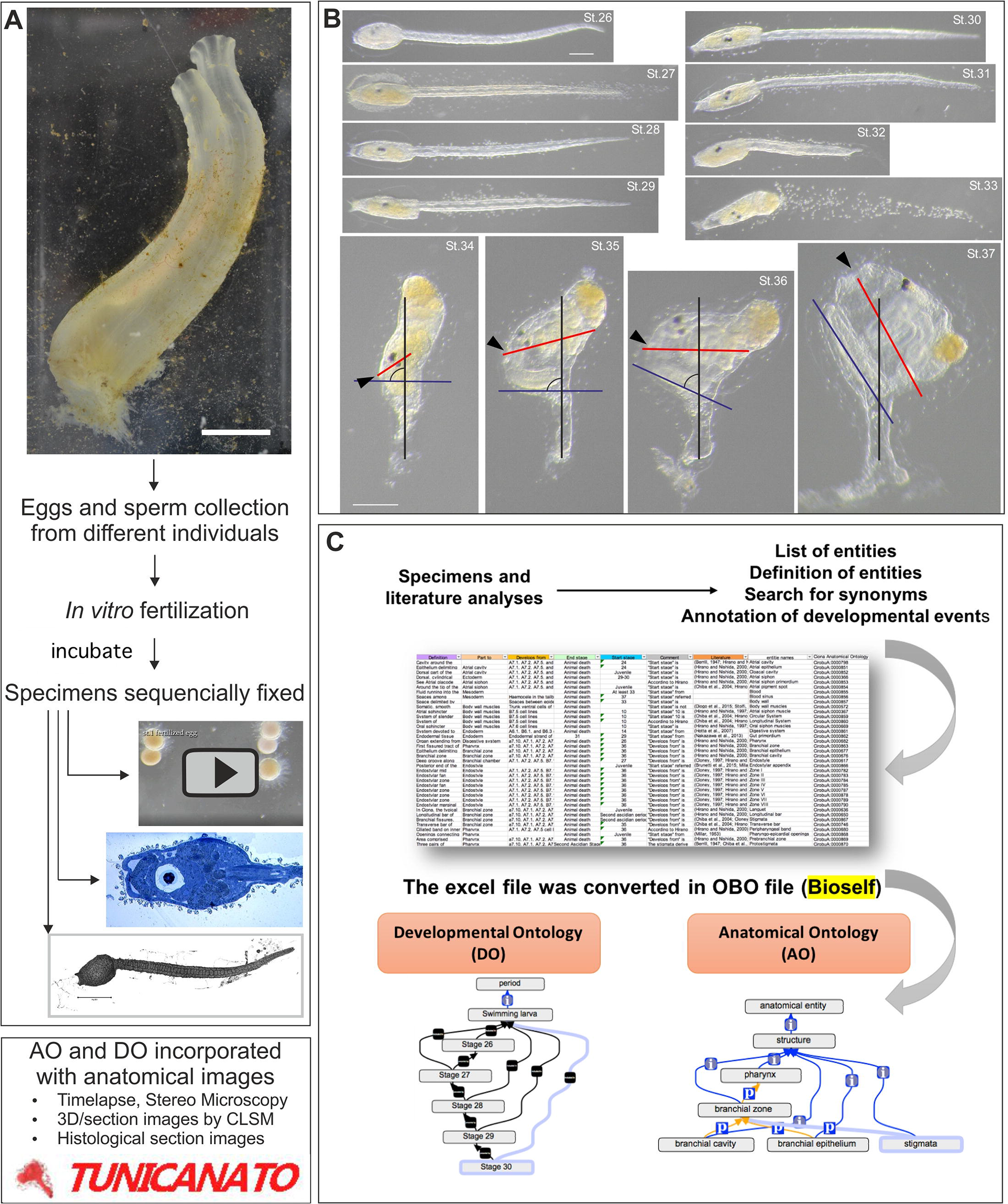
Methodological procedure to produce the ontology. **A.** Gametes were collected from adult individuals of *C. robusta* (*C. intestinalis* type A) for *in vitro* fertilization. At a specific time, samples were observed via stereomicroscope, photographed, and fixed for CLSM and histology. Scale bar 2 cm. **B.** Summary of Stages 26-37. Stereomicroscopy - *in vivo* specimens. In individuals belonging to Stages 34-36 (Body Axis Rotation Period), the blue line indicates the longitudinal body axes (the antero-posterior axis) that is parallel to the endostyle; the red line indicates the oral siphon-gut axis; the black line indicates the stalk axis. In an individual at Stage 37 (Juvenile Period), the rotation is almost completed and the longitudinal body axis is almost parallel to the stalk axis. Stages 26-33: anterior at left, left view; Stages 34-37: oral siphon (anterior) indicated by arrowhead; left view. Scale bar 100 μm. **C.** After definition of the developmental stages, specimens and literature were analyzed. After that, entities in hierarchical order, definitions, synonyms, developmental information, and literature were annotated in an Excel file. These data were edited using OBO-Edit (http://oboedit.org/), allowing for the visualization of relationships among entities. **D.** Lastly, data were associated with images and movies in the web-based resource TunicAnatO (https://www.bpni.bio.keio.ac.jp/tunicanato/3.0/).

From now on, each entity, both developmental and anatomical, is written in bold when introduced for the first time; relations between entities appear in italics, while entity definitions appear between quotation marks. In the ontology, an identification (ID) code has been assigned to each anatomical and developmental entity. ID, which here is in brackets, is a set of numbers preceded by two prefixes. The first prefix is “Cirobu”, referred to the species name *<u>C.</u> <u>robu</u>sta* (*C. intestinalis* type A). The second prefix follows the first one and is “A” when the ID is referring to the anatomy and “D” when it is referring to development.

The larva stages considered here belong to the **Larva Period** (CirobuD:0000013), which is included in the **Embryonic Development, Pre-Metamorphosis Meta-Period** (CirobuD:0000003). The **Metamorphosis Meta-Period** (CirobuD:0000004) is divided into the following Periods: **Adhesion** (stage 30; CirobuD:0000013), **Tail Absorption** (stages 31-33; CirobuD:0000015), and **Body Axis Rotation** (stages 34-36; CirobuD:0000016). The **Post-Metamorphosis Meta-Period** (CirobuD:0000005) consists of the **Juvenile Period** (CirobuD:0000017). Overall, 41 stages (12 of which here described and integrated with original images) until the Juvenile Period were defined and combined with the previous ontology ^24^ (Supplementary Table S3).

Once we defined the DO, we constructed the AO (Fig. 1). We carefully studied our anatomical data, comparing stage-by-stage information from CLSM and histology. This allowed us to recognize all the organs/tissues and follow their differentiation over time. We then listed, in an Excel file, the terms referring to the recognized anatomical entities (including synonyms, when present) reported in the literature and used by researchers since 1893 (Supplementary Data S6). We listed 203 entities (Supplementary Data S7, column F, “Further specification 4”), assigning to each one its ID. Moreover, we detailed, for each anatomical entity, the following characteristics: the definition (column “Definition” in Supplementary Data S7, Supplementary Data S8) based on the literature; the anatomical hierarchical level, specifying to which superior entity each belongs (*Part of*); the tissue from which it derives (*Develops from*); the developmental stage of its disappearance (*End stage*); and the developmental stage in which it is first recognizable (*Start stage*). Therefore, the *Start stage* and *End stage* relationships link the AO to the DO, providing the precise description of the timing of development. If necessary, we took note of a specific feature (column Comment in Supplementary Data S7) and listed the bibliographic references (Supplementary Data S6 and column Literature in Supplementary Data S7). We also built a complete anatomical atlas in which most of the anatomical entities were efficiently annotated (Figs. 2–5; Supplementary Figures S9 - S15). Lastly, all the curated data were incorporated into a computable OBO format ^34^ (Fig. 1).

**Figure 2.**
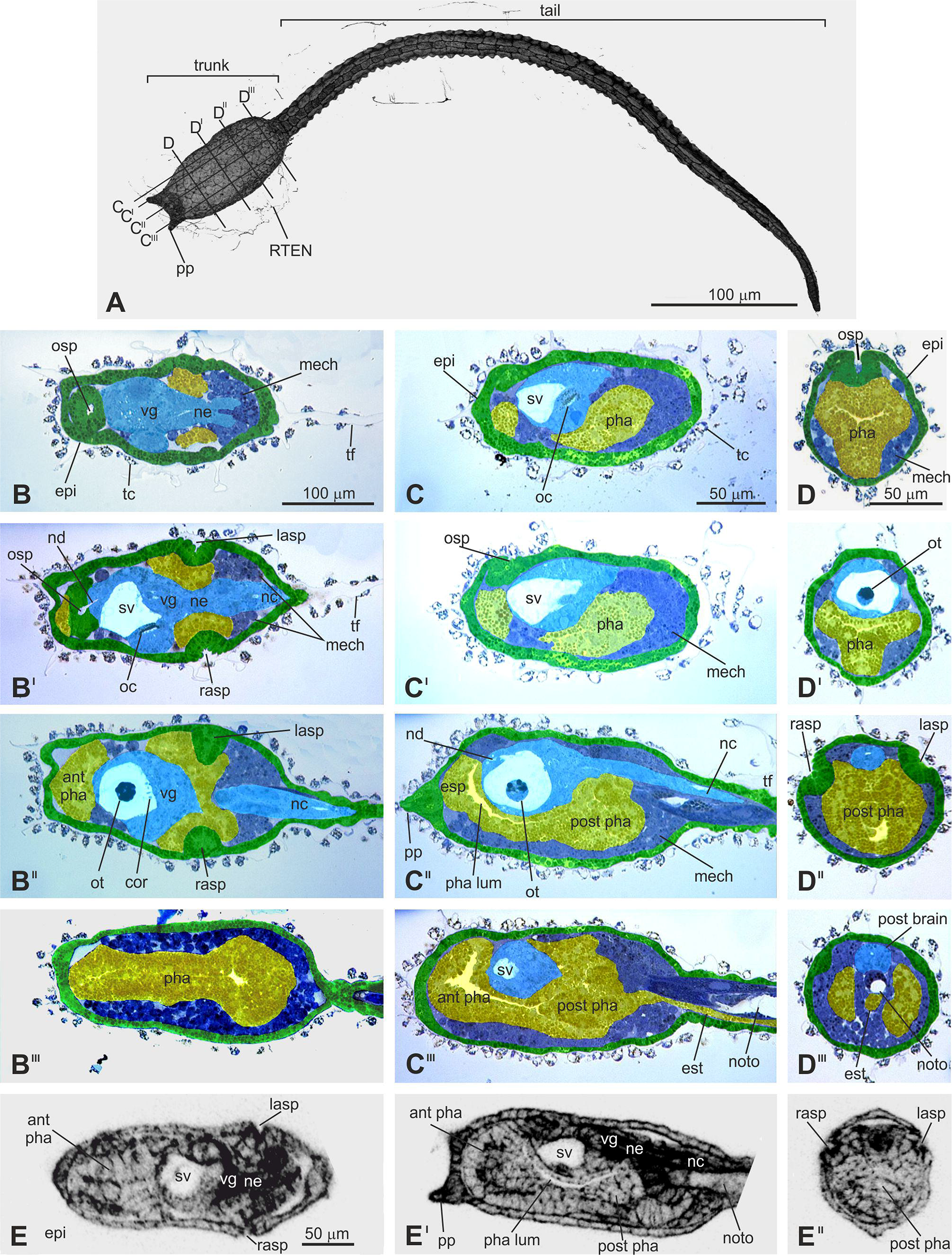
Early swimming larva (Stage 27). **A**. Larva, dorsal view. Lines on the larval trunk labeled by C-C^III^ and D-D^III^ indicate levels of sagittal and transverse sections shown in C-C^III^ and D-D^III^, respectively. CLSM. **B-D**^**III**^. Selected sections from complete datasets of serially sectioned larvae. B-B^III^ frontal sections from the dorsal to ventral sides; C-C^III^ sagittal sections from the right to left sides; D-D^III^ transverse sections from the anterior to posterior sides. Light microscopy, Toluidine blue. Enlargements in B^I^-B^III^, C^I^-C^III^, and D^I^-D^III^ are the same as in B, C, and D, respectively. Green: ectodermal non-neural tissues; yellow: endodermal tissues; light blue: ectodermal neural tissues; dark blue: mesodermal tissues. **E-E**^**II**^. Frontal (E), sagittal (E^I^), and transverse (E^II^) optic sections of the same larva. CLSM. Enlargement is the same in E-E^II^. For used symbols, see B-D^III^. ant pha: anterior pharynx; epi: epidermis; esp: endostyle primordium; est: endodermal strand; lasp: left atrial siphon primordium; mech: mesenchyme; nc: nerve cord; nd: neurohypophyseal duct; ne: neck; noto: notochord; oc: ocellus; osp: oral siphon primordium; ot: otolith; pha: pharynx; post pha: posterior pharynx; pp: papilla; rasp: right atrial siphon primordium; RTEN: cilium of a rostral trunk epidermal neuron; sv: sensory vesicle; tc: test cell; tf: tail fin; vg: visceral ganglion.

For example, for the entity **larval central nervous system** (CirobuA:0000579), the AO provides a consistent classification of cell types, tissues, and structures. Its relationship to the upper-level term **larval nervous system** (CirobuA:0000658) indicates that the larval central nervous system is *part of* the latter. The AO shows that the organ at stage 22 *develops from* its precursors, the A8.7, A8.8, A8.16, a8.17, a8.18, a8.19, a8.25, and b8.19 cell lines. In this example, the developmental relation is based on data from the cell lineage ^35–37^. The larval central nervous system regresses (*End stage*) at stage 33 (Stage Late Tail Absorption) when most of the larval structures are reabsorbed at metamorphosis. The forebrain (*i.e.*, the anterior **sensory vesicle**, CirobuA:0000357), the midbrain (*i.e.*, the **neck**, CirobuA:0000657), and the hindbrain (*i.e.*, the **visceral ganglion**, CirobuA:0000778), are part of the **brain** (CirobuA:0000582). The forebrain contains 16 distinct entities, whereas the midbrain contains nine entities. The larval central nervous system includes four entities: **sensory vesicles** (CirobuA:0000938), **neck** (CirobuA:0000657), **visceral ganglion** (CirobuA:0000778), and **tail nerve cord** (CirobuA:0000740). Some of them include further sub-structures (“Further specification” columns in Supplementary Data S7). For example, within the **sensory vesicle** (CirobuA:0000938), five further entities are included (anterior sensory vesicle, **posterior sensory vesicle** (CirobuA:0000696), **coronet cells** (CirobuA:0000890), **ocellus** (CirobuA:0000666), and **otolith** (CirobuA:0000671)), whereas the visceral ganglion includes the **motor neurons** (CirobuA:0000891). In addition, ependymal cells are included in the anterior sensory vesicles, neck, visceral ganglion, and tail nerve cord. For example, in the visceral ganglion, they are the lateral (CirobuA:0000643), ventral (CirobuA:0000776), and dorsal (CirobuA:0000606) visceral ganglion ependymal cells, respectively.

Below, we first describe how, where, and when the complex anatomical structures of *C. robusta* (*C. intestinalis* type A) emerge and change during the Periods defined in this study (see Supplementary Data S2 for detailed description of stages). Then we present an overview of the number of anatomical entities and their appearance throughout the entirety of ontogenesis.

### The Embryonic Development, Pre-Metamorphosis Meta-Period (Stages 26 to 29)

For the Embryonic Development, Pre-Metamorphosis Meta-Period, we described the last Period, called the Swimming Larva Period, during which the hatched larva (17.5hpf) swims actively, beating its tail. Although larvae belonging to this Period are generally defined as “swimming” larvae, their internal structures change significantly over time. Therefore, the Period (17.5-24 h after fertilization at 18°C) was divided into four anatomically distinguishable Stages, from Stage 26 to Stage 29, until the end of the locomotion phase (Fig. 2; Supplementary Figure S9-S11).

Up to 90 entities are histologically recognizable as larval organs (Supplementary Data S7). The main larval trunk territories are: the epidermis (CirobuA:0000619); the **endoderm** (CirobuA:0000615); the **mesenchyme** (CirobuA:0000653), mainly in the ventral-lateral trunk (**trunk ventral cells** (B7.5 line): CirobuA:0000748); and the nervous structures, such as the sensory vesicle, the ocellus, and the otolith, in the dorsal-mid trunk. The **nerve cord** (CirobuA:0000740), the **notochord** (CirobuA:0000665), the **muscles** (CirobuA:0000739), and the **endodermal strand** (CirobuA:0000616) are in the tail. As is typical in ascidians, the larva possesses three **adhesive papillae** (CirobuA:0000675) on the anterior trunk tip: two dorsal and one ventral.

### Metamorphosis Meta-Period (24-60 h at 18°C, Stages 30 to 36)

In this Meta-Period, which is triggered by the larva adhesion, several anatomical changes are simultaneously observed, as the larval organs degenerate, whereas the adult organs appear and differentiate (Figs. 3–5; Supplementary Figures S12-S14). It is subdivided in three Periods: Adhesion, Tail Absorption, and Body Axes Rotation.

**Figure 3.**
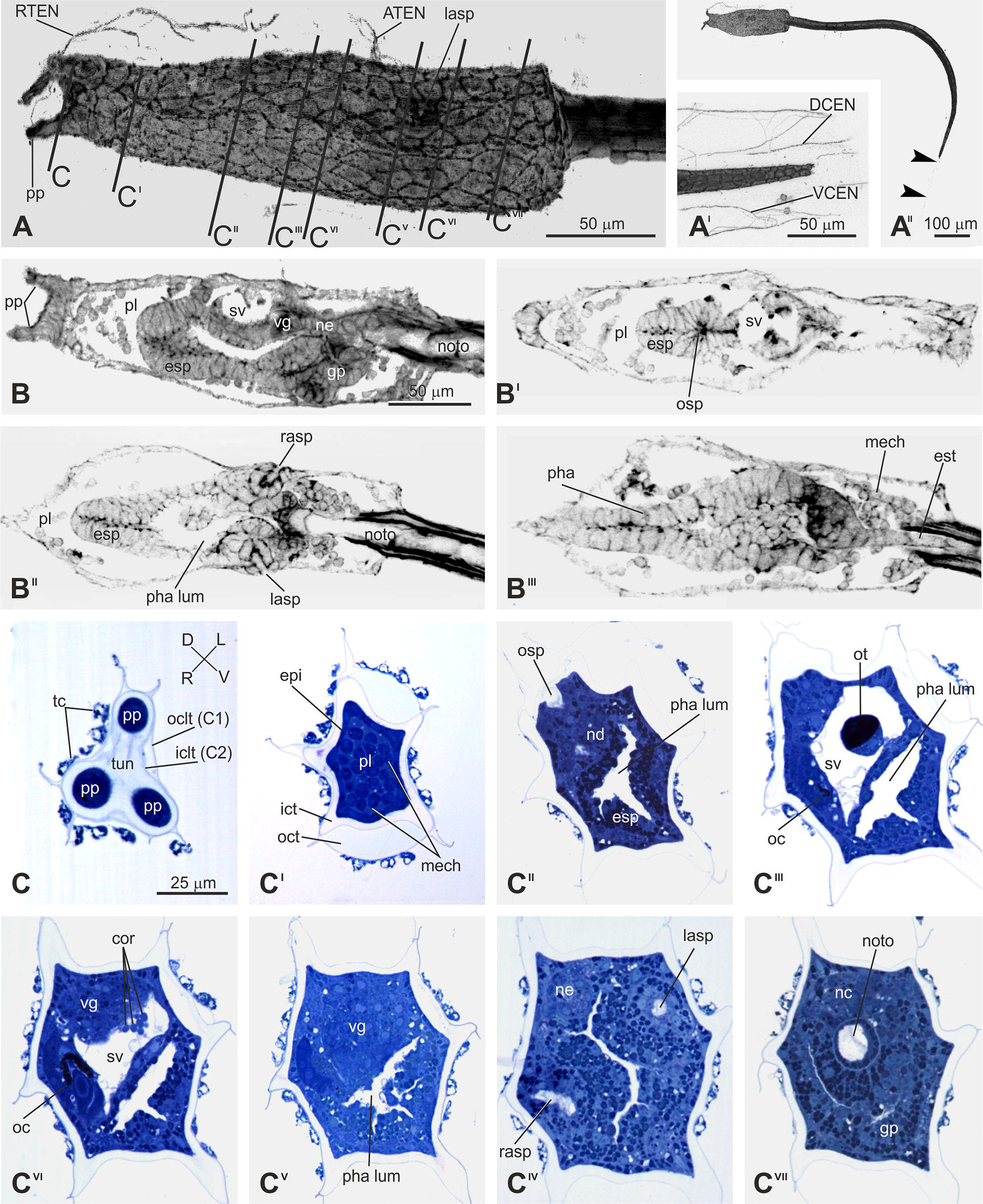
Adhesion (Stage 30). **A-A**^**II**^. Trunk (A), tail tip (A^I^), and larva (A^II^) in adhesion, seen from the left side. CLSM. Lines on the larval trunk labeled by C-C^VII^ indicate levels of cross-sections shown in C-C^VII^. Arrowheads in A^II^: tunic remnant at the tail tip. Enlargement is the same in A-A^II^. **B-B**^**III**^. A medial sagittal (B) and three frontal (from dorsal to ventral side) (B^I^-B^III^) optic sections of the larval trunk. CLSM. Enlargement is the same in B-B^III^. **C-C**^**VII**^. Eight transverse sections of the same larva (from anterior to posterior side). Light microscopy, Toluidine blue. ATEN, DCEN, RTEN, VCEN: cilium of an anterior trunk, dorsal caudal, rostral trunk, and ventral caudal epidermal neuron, respectively; cor: coronet cells; epi: epidermis; esp: endostyle primordium; esr: endodermal strand; gp: gut primordium; iclt (C2) and oclt (C1): inner (C2) and outer (C1) cuticular layer of the tunic, respectively; ict and oct: inner and outer compartment of the tunic, respectively; lasp: left atrial siphon primordium; mech: mesenchyme; nc: nerve cord; nd: neurohypophyseal duct; ne: neck; noto: notochord; oc: ocellus; ot: otolith; pha lum: pharynx lumen; pl: preoral lobe; pp: ventral papilla; rasp: right atrial siphon primordium; sv: sensory vesicle; tc: test cell; tf: tail fin; vg: visceral ganglion.

### The Adhesion Period

The Adhesion Period (24-27 h post-fertilization at 18°C) consists of one stage: Stage 30 (Fig. 3). The Period triggers the metamorphosis, starting important anatomical and developmental modifications in the larva. The latter stops swimming and attaches to a suitable substrate through its adhesive papilla. The adhesive larval papillae retract during this period.

### The Tail Absorption Period

Immediately after the adhesion, ascidian tadpole larvae lose their tail by tail regression during tail absorption period. The first observable change in the initiation of *Ciona* metamorphosis, earlier than the onset of tail regression, is the backward movement of the posterior trunk epidermis^38^ thereafter the larval tail (**absorbed larval tail**: CirobuA:0000951) begins to be absorbed (27-30 h post-fertilization at 18°C). This Period consists of three stages: Stage 31, Stage 32, and Stage 33, whose duration depends on the extent of tail regression. Usually the Tail absorption is completed in 75 - 90 min^39^(Supplementary Video 3; Supplementary Figures S12-S13) at 18 - 20°C. At 20°C, the earliest tail absorption start time was 6.3 hour after hatching (hour post hatch: hph), the latest time was 17.9 hph, and the average time was 9.9 hph (N = 37). The earliest time for tail absorption to 50% tail length was 6.5 hph, the latest time was 18.3 hph, and the average time was 10.4 hph (N=43). The earliest time for tail absorption was 7.9 hph, the latest time was 18.8 hph, and the average time was 11.4 hph (N=43). From the above, it takes about 30 minutes from the beginning of tail absorption to 50% of tail absorption, and about 1 hour from the tail absorption of 50% to completion of tail absorption (Table 1).

### The Body Axis Rotation Period

After the tail absorption, the **Body Axis Rotation Period** (CirobuD:0000016; 30-60 h post-fertilization at 18°C) occurs. Ascidian metamorphosis is characterized by the rotation of inner organs through an arc of about 90° ^30^. The adult ascidian has a longitudinal body axis (the antero-posterior axis) that is parallel to the endostyle and passes through the oral siphon and the gut. In the adhering larva, the longitudinal body axis, easily individuated by the endostyle, is parallel to the substrate. During metamorphosis, it rotates progressively so that, at the end of metamorphosis, it is almost perpendicular to the substrate and aligned with the stalk. This Period consists of three stages: Stages 34, 35, and 36 (Fig. 4; Supplementary Figure S14; Fig. 5), depending on changes in body shape (in particular, the enlargement of the branchial chamber due to protostigmata perforation) and the angle formed by the stalk axis (the definitive longitudinal body axes) and the endostyle axis. The latter, in early metamorphosis, does not correspond precisely with the oral siphon-gut axis, as the tail remnants occupy a large posterior body portion.

**Figure 4.**
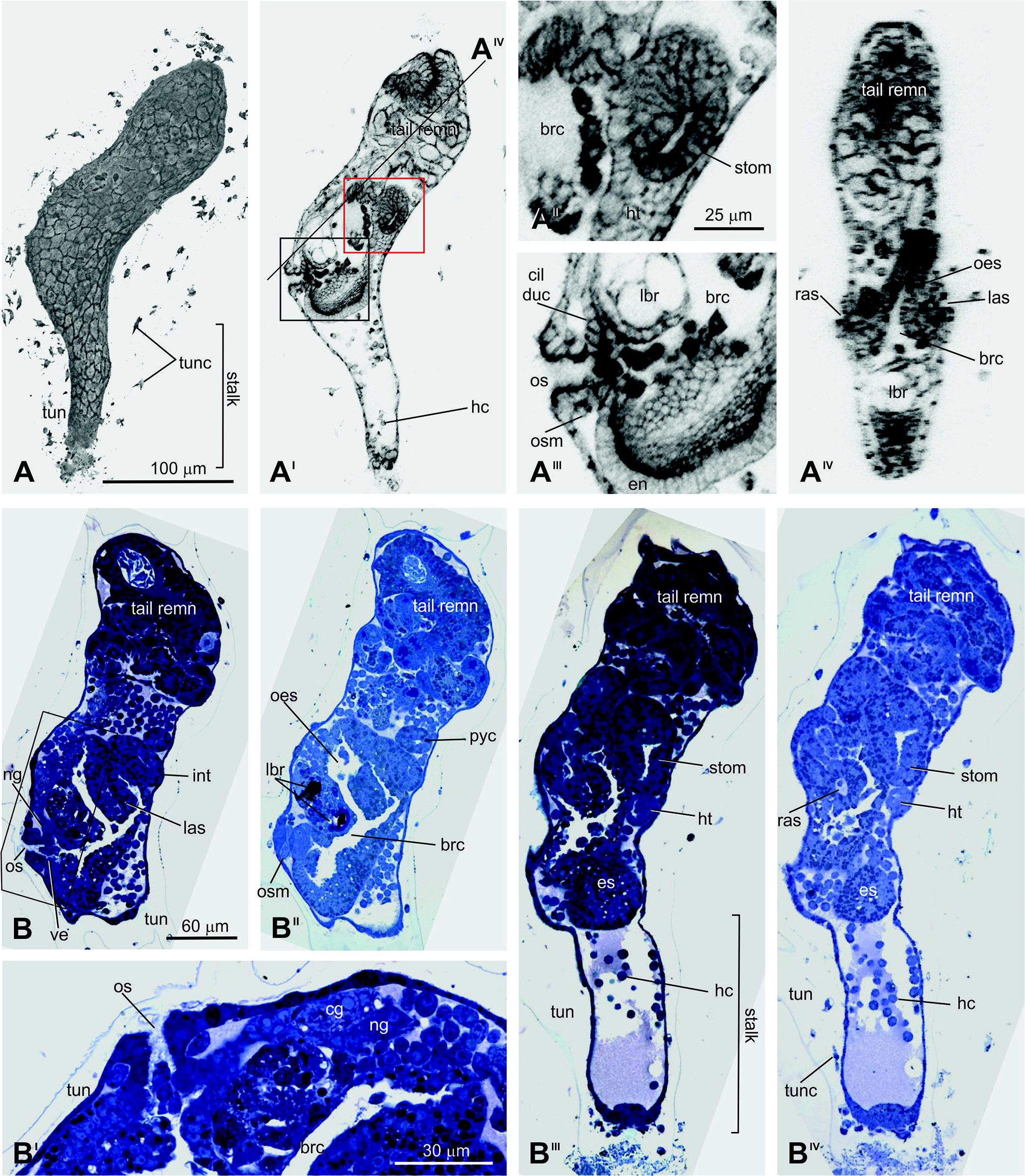
Early body axis rotation (Stage 34). **A-A**^**IV**^. Metamorphosing larva seen from the left side (A) and its medial sagittal optic section (A^I^). In A^I^, the area bordered by the red line is enlarged in A^II^; that one bordered by the black line is enlarged in A^III^; the line marked by A^IV^ represents the level of section A^IV^. Enlargement is the same in A-A^I^, and in A^II^-A^IV^. CLSM. **B-B**^**IV**^. Serial sagittal histological sections of a metamorphosing larva from the left to right sides (B, B^II^-B^IV^); the area bordered by the black line in B is enlarged in B^I^ to show the oral siphon and neural complex. Enlargement is the same in B and B^II^-B^IV^. Toluidine blue. brc: branchial chamber; cil duc: ciliated duct of the neural gland; cg: cerebral ganglion; es: endostyle; hc: maemocytes; ht: heart; las: left atrial siphon; int: intestine; lbr: larval brain remnants; ng: neural gland; oes: oesophagus; os; oral siphon; osm: oral siphon muscle; pyc: pyloric caecum; ras: right atrial siphon; stom: stomach; tail remn: tail remnants; tun: tunic; tunc: tunic cells.

**Figure 5.**
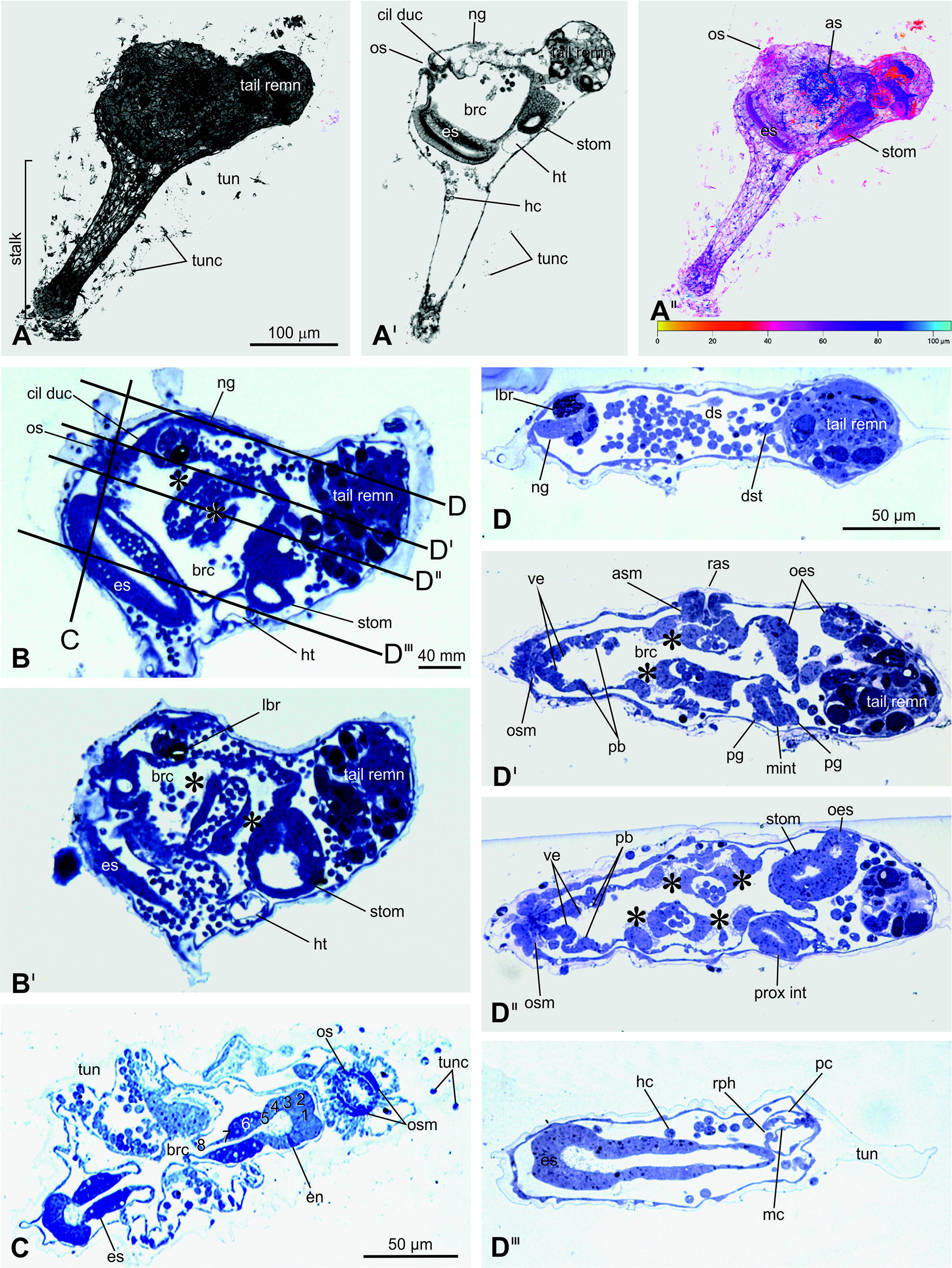
Late body axis rotation (Stage 36). **A-A**^**II**^. Metamorphosing larva, seen from the left side (A), its medial sagittal optic section (A^I^), and its depth-coded image (A^II^). The depth information is represented by a heat map: warmer colors go to the front, and cooler colors to the back. Color bar: value of depth (μm). CLSM. Enlargement is the same in A-A^II^. **B-B**^**I**^. Two sagittal sections of a metamorphosing larva. In B, lines on the larval trunk, labeled by C and D-D^III^, indicate levels of the transverse section and the frontal sections shown in C and D-D^III^, respectively. Asterisks: protostigmata. Toluidine blue. Enlargement is the same in B-B^I^. **C.** Transverse section of the metamorphosing larva at endostyle and oral siphon level. Note the differentiating eight zones in endostyle (1-8). Toluidine blue. **D-D**^**III**^. Serial frontal histological sections of a metamorphosing larva from the dorsal (D) to ventral (D^III^) sides. Toluidine blue. Enlargement is the same in D-D^III^. brc: branchial chamber; cil duc: ciliated duct of neural gland; ds: dorsal sinus; dst: dorsal strand; es: endostyle; hc: haemocytes; ht: heart; lbr: larval brain remnant; mi: medium intestine; mc: myocardium; ng: neural gland; oes: oesophagus; os: oral siphon; osm: oral siphon muscle; pb: peripharyngeal band; pc: pericardium; pg: pyloric gland; prox int: proximal intestine; rph: raphe; rpsm: right protostigmata; stom: stomach; tail remn: tail remnants; tun: runic; tunc: tunic cells; ve: velum.

During this Period, many tunic cells in the tunic actively change their shape, forming filopodia, indicating that these cells are mobile and differentiated (“tunc” in Supplementary Figure S14A and in Fig. 5A).

### The Postmetamorphosis Meta-Period (3 days to over 7 days at 18°C, stages 37 to 43)

The **Postmetamorphosis Meta-Period** (CirobuD:0000005) consists of three Periods: the **Juvenile Period** (CirobuD:0000017), the **Young Adult Period** (CirobuD:0000018), and the **Mature Adult Period** (CirobuD:0000019). The Juvenile Period (3 days to over 7 days post-fertilization at 18°C) consists of five stages: 37, 38, 39, 40, and 41 (Supplementary Table S3 and Supplementary Figure S15), defined mainly by gill slit and gut elaboration. Individuals still do not have mature reproductive organs, although gonads are developing. As reported above, this ontology includes only the description of Stage 37. The Young Adult Period consists of Stage 42 (2nd Ascidian Stage), corresponding to Stage 8 in Chiba et. al., 2004.

### Number of anatomical entities and their appearance during ontogenesis

All the anatomical entities annotated in the ontology, from both present results and previously reported data ^24^, were analyzed in whole during the complete ontogenesis of *Ciona*. Figure 6 presents, stage by stage, their number from Stage 0 (unfertilized egg) to animal death. The entities associated to the embryo/larval life is 88/203 (Fig. 6, yellow column), entities associated to the juvenile/adult life is 93/203 (Fig. 6, red column) and entities persistent in biphasic life is 22/203 (Fig. 6, blue column). The graph shows that there are two rounds of tissue/organ increase. The first one is marked and occurs after fertilization; it reaches maximum number at Stage 25 (Mid-gastrula), when about 141 entities are recognizable. At Stage 34 (corresponding to the conclusion of the Body Axis Rotation Period), there is a sharp decrease in the number of entities, which drops to 90. This decrease is followed by a second increase that reaches 114 entities to the Juvenile Period.

**Figure 6.**
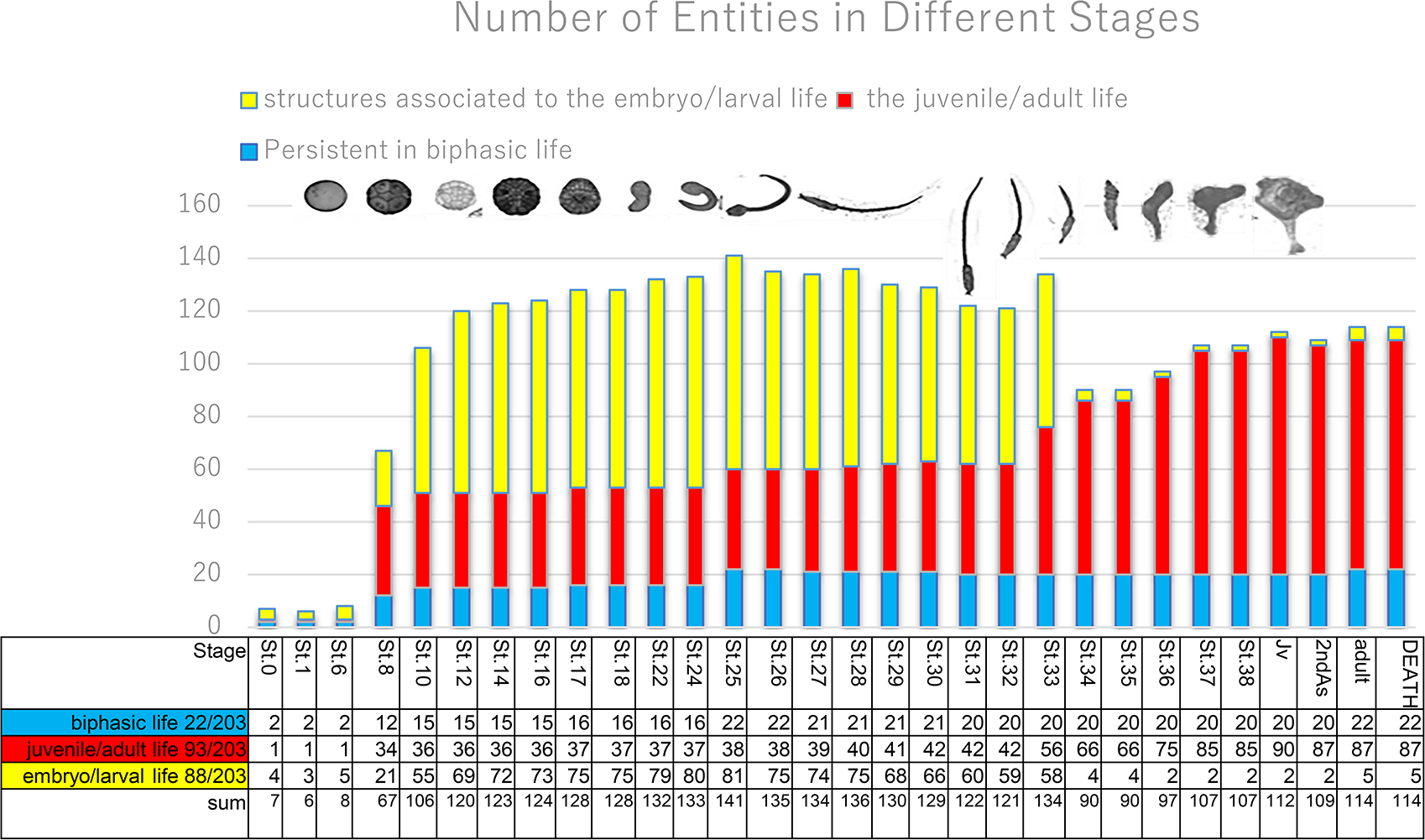
Number variation of anatomical entities in each developmental stage. Graph and matrix showing the variation in number of anatomical entities during development (Stages in abscissa), from Stage 0 (unfertilized egg) to animal death. Yellow columns refer to cellular or structural level entities associated to the embryo/larval life (88/203), red columns refer to cellular or structural level entities associated to the juvenile/adult life (93/203) and blue columns refer to cellular or structural level entities (22/203) persistent in both biphasic life. Jv: Juvenile Period; 2^nd^: 2^nd^ Ascidian Stage; adult: Adult Stage; Death: Animal Death.

## Discussion

### The ontology of *Ciona*: a powerful tool for developmental biology studies

*Ciona* is considered a valuable model for studying the developmental biology of tunicates and the evolution of chordates. For this species, several databases have contributed as resources for genome and gene expression information (*e.g.*, Ghost, http://ghost.zool.kyoto-u.ac.jp/indexr1.html; CITRES, https://marinebio.nbrp.jp/ciona/) and proteomic studies (*e.g*., CIPRO, http://cipro.ibio.jp/), or provide technical information (*e.g*., ACBD, the Ascidians Chemical Biological Database, https://www.bpni.bio.keio.ac.jp/chordate/acbd/top.html). All the available databases are easily consultable via the Tunicate Web Portal (https://tunicate-portal.org/). In this study, we present the database TunicAnatO, which implements the previous database FABA ^24^ and is devoted to anatomy and development. Moreover, the ADO built here is combined with the ANISEED database (https://www.aniseed.cnrs.fr), which provides high-throughput data and *in situ* experiment data from the literature for ascidian species. Therefore, we integrate the panorama of actual databases and offer a tool that will help researchers in the recognition the anatomical structures of their interests. This will allow for the standardization of data underpinning an accurate annotation of gene expression and the comprehension of mechanisms of differentiation.

### The developmental ontology

In this work, merging the previously reported developmental stages ^24,33^, with new data from stereomicroscopy, CLSM, and histology, we implemented the description of the whole life cycle of *Ciona*, from fertilization to juvenile. The whole development has been divided into Meta-Periods, Periods, and Stages, following the canonical temporal subdivision of developmental ontologies ^33^. Using a low-resolution microscope for dissection to examine larvae, metamorphosing individuals, and juveniles, we defined the new subdivision into stages. The simplicity of stage recognition is a prerequisite for a good staging method. Researchers will easily be able to discriminate stages, using a simple instrument, a stereomicroscope, when checking the development of their living samples in the laboratory after *in vitro* fertilization, or when analyzing fixed whole-mount specimens.

We introduced 12 new stages (from Stage 26, Hatching Larva, to Stage 37, Early Juvenile I) that add to those already reported up to the Larva Stage ^24^. Therefore, 37 stages are now described in detail and documented with original images. Considering that we also defined (without describing) Stage 38 (Early Juvenile II) and Stage 39 (Mid Juvenile I), once they are described, the whole Juvenile Period will be completed. The lacking steps are, then, the Young Adult Period (Stage 40) and the Mature Adult Period (Stage 41). However, for the latter Period, the exhaustive anatomical description by Millar ^32^ is still an essential reference. In summary, the whole life cycle of *Ciona* is almost described and annotated. This is an important result, considering that ontologies regarding other model organisms are limited to embryogenesis ^1,3–5,40^.

It should be noted that, in annotating the progressive organ appearance and degeneration, we could also describe in detail the metamorphosis process, whose general reports for ascidians are dated, not so accurate and timed, and limited to a few species (see for review: ^29,41^). Only some specific processes occurring during metamorphoses, such as tail regression ^39,42^ or papillae retraction ^42^, have been described in detail in *C. robusta* (*C. intestinalis* type A).

### The anatomical ontology

This study underlines the importance of a combined analysis of data. In fact, for each stage, we examined corresponding high-resolution images thanks to CLSM and histology. The two methods have advantages and disadvantages in terms of studying anatomy. CLSM provides high tissue resolution and relatively rapid processing, so it allows for the analysis of multiple samples. Moreover, in automatically making z-stacks, we can quickly obtain 3D reconstructions. However, tissues/organs can be difficult to recognize due to the limited number of fluorochromes that can be simultaneously used. Moreover, a low laser penetration can be a limit for the study of thick specimens. On the other hand, histology offers (other than a high tissue resolution) an easy tissue/organ recognition thanks to the different tissue affinities to labeling. However, the method is time-consuming, which means that few samples can be analyzed, and 3D reconstructions are not automatically generated. Therefore, we used, in combination, the complementary information coming from these two working methods, making it possible to identify, with precision, the inner structures as well as the outer surface of individuals, annotating in total 203 anatomical territories.

To build the hierarchical tree of anatomical entities (specified by the relation *Part of* in Supplementary Data S7) and to define each of them (complete with synonyms), we consulted several publications covering over 120 years of literature on *Ciona* and other ascidians, from 1893^31^ to today. Fundamental references for creating the dictionary were, among others, those published by Millar ^32^, Kott ^43^, Burighel and Cloney ^41^, Chiba and collaborators ^33^, and the last description of the species by Brunetti and collaborators^21^. We also consulted the glossary TaxaGloss (https://stricollections.org/portal/index.php).

The AO is documented by the database TunicAnatO, which is an anatomical atlas of original images readily available via the internet and easily accessible from any standard web browser (https://www.bpni.bio.keio.ac.jp/tunicanato/3.0/). This database contains information from both z-slice sections and 3D reconstruction images, and histological sections at each time point along the developmental course of *C. robusta* (*C. intestinalis* type A). In images, the anatomical entities were labeled, providing a guide for tissue recognition.

### Data integration in anatomical and developmental ontology

In this work, we linked DO and AO in a comprehensive DAO, as we defined, for each entity, the relations *Develops from*, *Start stage*, and *End stage*. These relations were determined at the cell level when cell lineage data were available and at the tissue/organ level where complexity did not allow for the following of cell genealogy. Lastly, when possible, we also annotated features linked to organ functionality (swimming, feeding, respiration, or heart beating). Some entities showed multiple possibilities to be defined, while others had uncertain/controversial definitions.

In some cases, and where possible, we defined the anatomical entities according to multiple organizational levels. For example, the “atrial siphon muscle” *Start stage* is Stage 10 if we refer to cell lineage ^44–46^, while it is Stage 33 if we refer to histology (muscles recognizable on sections) and Stage 36 if we refer to the functional state of the muscles (ability to contract). Similarly, the *Start stage* for the “endostyle” is Stage 27 if we refer to the cell lineage ^47^, while it is Stage 34 if we consider the presence of its main histological features (its subdivision into eight symmetrical zones, visible on sections) and Stage 36 if we refer to its physiological activity during feeding (mucus production trapping food particles). These and other, similar examples, exhibiting multiple tissue recognition levels, are all annotated in the ontology (among the Comments in Supplementary Data S7), for a comprehensive view of development. It is to note that in *Ciona* the cell lineage is not known for a few anatomical entities. In these cases (referred mainly to larval pharynx, tail epidermis and some larval brain components), we referred to data from *Halocynthia roretzi* ^35,47,48^, specifying this in the Comments sections of the Supplementary Data S7. This information represents an important reference for future studies on *Ciona* development. The above-reported examples highlight the complexity and choices underlying the ontology building. However, the presence of comments associated with each entity in this ontology, and the huge number of citations reported, assure users of a comprehensive view of *Ciona* anatomy and development.

An analysis of other ontologies currently available shows that the ontology of *Ciona* presented here is very rich in information. Sixty-one ontologies deal with anatomy on FAIR sharing (https://tinyurl.com/ybhhfd8c)^49^. Among them, 12 describe the anatomy of animal model organisms (*e.g*., *Drosophila*, *Caenorabditis elengans*, mosquito, mouse, zebrafish, *Xenopus*, planaria, the ascidian *Botryllus schlosseri*). In terms of a comparison of the ontology of *C. robusta* (*C. intestinalis* type A) presented here with the latter (Supplementary Table S16), the first one possesses very rich lineage information compared to other ones. Moreover, among the 12 above-mentioned ontologies, four combine developmental stages and anatomical terms, eight include the relation *Develops from*, and 10 use references as a source of data. The ontology of *Ciona* exhibits all these features. It is to be considered, however, that ontologies are never-ending tools. They must be continuously updated when new information becomes available.

### An overview of the ontogenesis of *C. robusta* (*C. intestinalis* type A)

Thanks to the annotation of the relations, *Start stage* and *End stage*, we could verify, in *Ciona*, the progressive emergence—and, where appropriate, disappearance—of its unique features. Looking at them as a whole, we obtained a global view of ontogenesis.

Our results show that ascidians have two rounds of increasing complexity: the first one during cleavage until gastrulation, and the second one during metamorphosis. This can reflect the development of structures associated with the larval life (88 in total; for example, the larval nervous system, the tail with associated notochord and muscles) and with the juvenile/adult life (93 in total; for example, the branchial basket, the gonad, the gut). Other structures (22 in total) are, of course, persistent throughout the whole ontology: They are, for example, the epidermis, the tunic, and the hemocytes.

Moreover, we show that almost one-third of the anatomical structures disappear from stage 33 (134 entities) to stage 34 (90 entities) (Fig. 6). This occurs during the Tail Absorption Period and the beginning of the Body Axis Rotation Period, when structures exclusively formed for the larval life degenerate. This drastic event was not previously documented quantitatively. It should also occur in other invertebrate species, such as barnacles and sea urchins ^50,51^. In fact, there is the loss of many organs associated with the motile larva that metamorphoses in a stationary form. Such a sharp decrease in anatomical structures was not reported in other chordate animals.

## Conclusions

In this study, we present the ADO of the ascidian *C. robusta* (*C. intestinalis* type A), from the swimming larva stage, through metamorphosis and until the juvenile stages. We define 12 stages that, together with the previously described stages related to embryogenesis, extend our knowledge to almost the whole ontogenesis. This ontology, providing the hierarchical description of more than 203 anatomical entities, complete with definitions, synonyms, and bibliographic references, provides the guideline for several functional studies on tunicate cell biology, development, and evolution. It allows for the standardization of data underpinning the accurate annotation of gene expression and the comprehension of mechanisms of differentiation. It will help in understanding the emergence of elaborated structures during both embryogenesis and metamorphosis, shedding light on tissue degeneration and differentiation occurring at metamorphosis.

## Methods

### Biological materials

*C. intestinalis* type A (*C. robusta*) adults for time-lapse imaging and for confocal scanning laser microscopy (CLSM) were provided by NBRP from the Maizuru bay and Tokyo bay areas in Japan. For histology, adults were obtained from the Lagoon of Venice, Italy. Species determination was performed checking the discriminating factor “trunk shape” of late larvae, described in Pennati et al., 2015. Specimens collected in different sites possessed the same anatomical and developmental features.

### Preparation of embryos for time-lapse imaging

Eggs and sperm were obtained surgically from gonoducts. After insemination, eggs were maintained in agarose-coated dishes with Millipore-filtered seawater (MFSW) containing 50 μg/ml streptomycin sulfate, and the early cleavages were uniformly synchronized (data not shown). To keep the temperature stable, we used a Peltier-type incubator (CN-25B, Mitsubishi, Japan) without any vibration to prevent embryo fusion. Embryos developed in hatched larvae approximately 18 h after insemination.

The naturally hatched larvae derived from egg with chorion were maintained in plastic dishes on the thermo-plate at 20°C to acquire images. Using a digital camera (Olympus SP-350) mounted on a microscope, images were acquired every 3 to 10 min for 7 days (Supplementary Video S4). After 3 days post-fertilization, food was given (vegetal plankton, sun culture).

### Image acquisition at confocal scanning laser microscopy (CLSM)

Fixed specimens were prepared at different timings of development, from hatching larva up to 7dpf juvenile. Samples incubated at 18 ◻ were fixed for 30 min – 1 day at room temperature in 4% EM grade paraformaldehyde (nacalai tesque code 00380) in MOPS buffer (0.1 M 3-(N-Morpholino) propanesulfonic acid), adjusted to pH 7.5. Specimens were then washed three times with phosphate-buffered saline (PBS) and incubated in Alexa-546 phalloidin (Molecular Probes, Eugene, OR) in PBS containing 0.01% Triton X-100 (PBST) either overnight at 4°C or at room temperature for 1-2 h. Specimens were then rinsed for 3 min in PBS, attached to glass slide dishes, dehydrated through an isopropanol series, and finally cleared using Murray clear, a 2:1 mixture of benzyl benzoate and benzyl alcohol. Alexa 546 phalloidin was used to visualize cell membranes because it stains mainly cortical actin filaments.

Images were collected with a CLSM on a Zeiss LSM510 META with 40X oil objective or on an OLYMPUS fv1000. To reconstruct the 3D images, 100 cross-section images from top to bottom per sample were acquired (LSM image browser, Zeiss, Germany). The focus interval depended on the sample (from 0.5 to 1.2 μm). The resulting stacks were then exported to raw image series or to 3D image data for database integration. Although the timing of metamorphosis showed a huge deviation depending on the timing of adhesion, we considered an average timing, looking at animals exhibiting a representative morphology. Lastly, these stacks were integrated into the database TunicAnatO.

### Histology

After *in vitro* fertilization, larvae, metamorphosing individuals, and juveniles were fixed in 1.5% glutaraldehyde buffered with 0.2 M sodium cacodylate, pH 7.4, plus 1.6 % NaCl. After being washed in buffer and postfixated in 1% OsO_4_ in 0.2 M cacodylate buffer, specimens were dehydrated and embedded in Araldite. Sections (1 μm) were counterstained with Toluidine blue. Transverse, frontal, and sagittal serial sections were cut. Images were recorded with a digital camera (Leica DFC 480) mounted on a Leica DMR compound microscope. All photos were typeset in Corel Draw X5.

### AO/DO ID curation section

The anatomical and developmental terms with synonyms, definitions, and information about developmental events and anatomical entities were accumulated from textbooks, journals, and scientific observations. This information has been collected and formatted in two Excel files: one file on anatomy, the other on development. TunicAnatO was built in OBO format by using the open-source graphical ontology editor Open Biological and Biomedical Ontologies (OBO) edit ^34^.

## Supporting information

Supplementary_Figure_01_Life_Cycle

Supplementary_File_02_DescriptionStages2

Supplementary_File_03_Ciona_Staging_Table

Supplementary_File_06_Reference_List

Supplementary_File_07_Anatomical_Ontology (1)

Supplementary_File_08_DictionaryKH2

Supplementary_File_09_Stage_26

Supplementary_File_10_Stage_28

Supplementary_File_11_Stage_29

Supplementary_File_12_Stages_31-32

Supplementary_File_13_Stage_33

Supplementary_File_14_Stage_35

Supplementary_File_15_Stage_37

Supplementary_File_16_available_ontologies

## List of abbreviations

ADO: Anatomical Developmental Ontology
AO: Anatomical Ontology
CLSM: Confocal Scanning Laser Microscopy
DO: Developmental Ontology
ID: Identification
OBO: Open Biological and Biomedical Ontologies
TunicAnatO: Tunicate Anatomical and Developmental Ontology

## Declarations

### Ethics approval and consent to participate

Not applicable

### Consent for publication

Not applicable

### Availability of data and material

The datasets generated and/or analyzed during the current study are available in the following repositories, as well as our database, TunicAnatO (https://www.bpni.bio.keio.ac.jp/tunicanato/3.0/).

- Tunicate portal: https://tunicate-portal.org/resources/standards
- Bioportal: https://bioportal.bioontology.org/ontologies/CIROBUADO
- ANISEED: https://www.aniseed.cnrs.fr/aniseed/download/download_data
- FAIRsharing: https://fairsharing.org/bsg-s001475/

## Authors’ contributions

KH performed CLSM and time-lapse imaging, and constructed the website TunicAnatO. LM performed histology. KH and LM analyzed the microscopy images, described the development process, designed and built up the ontology, and drafted the paper. DD implemented entities into the OBO foundation. All participated in revising the manuscript. All authors read and approved the final manuscript.

## Acknowledgments

Authors thank Yutaka Satou Lab at Kyoto University, Manabu Yoshida Lab at the University of Tokyo and Onagawa Marine Station for providing individuals of *Ciona* samples with support by the National Bio-Resource Project of AMED, Japan; Akitsu Fukuzawa for collection of CLSM image stacks; Takumi T. Shito for the construction of the flash-independent version (version 3.0) of TunicAnatO; Paolo Burighel for the excellent comments and suggestions regarding the ontology.

## Supplementary Files

**Supplementary Figure S1**

File name: Supplementary_File_01_Life_Cycle.jpg

File format: .jpg

Title of data: Life cycle of *Ciona*

Description of data: Scheme of the life cycle of *C. robusta* (*C. intestinalis* type A). Blue box: developmental stages described in Hotta et al., 2007. Red box: developmental stages described in this paper.

**Supplementary Data S2**

File name: Supplementary_File_02_DescriptionStages.docx

Title of data: Description of features in each developmental stage. Each entity, both developmental and anatomical, is written in bold when introduced for the first time; relations between entities are in italics; entity definitions between quotation marks; ID in brackets. Abbreviations used in the cited Figures and Supplementary Figures are in italics in brackets.

**Supplementary Table S3**

File name: Supplementary_File_03_Ciona_Staging_Table

File format: .pdf

Title of data: Table of developmental stages in *Ciona*

Description of data: Table listing the Meta-Periods, Periods, and Stages of development of *Ciona*, from Stage 0 (unfertilized egg) to Stage 43 (adult). Main features, time of appearance after fertilization, and comparison with the staging method by Chiba and collaborators (Chiba et al., 2004) are reported for each stage. Yellow: stages defined in this work; Stages 26-37 are described in this work.

**Supplementary Video S4**

File name: Supplementary_File_04_Video

File format: .mov

Title of data: *C. robusta* (*C. intestinalis* type A) development

Description of data: Time-lapse movie showing the development of *Ciona* from the fertilized egg (Stage 1) to the juvenile (Stage 38). Observation was performed at 18°C for Stages 1-26 and at 20°C for Stages 27-37.

**Supplementary Figure S5**

File name: Supplementary_File_05_3D

File format: .tif

Title of data: Three-dimensional reconstructed images of *C. robusta* (*C. intestinalis* type A) post hatching stages

Description of data: Specimens labeled with Alexa 546 phalloidine (Molecular Probes). As the staining targets actin filaments, the cortical cytoplasm is stained in each cell. Stages 26-33: left view, anterio at left. Stages 34-37: leftview, anterior (oral siphon) at right (Stages 34-35) or top (Stages 36-37). Scale bar: 50 μm. CLSM.

**Supplementary Data S6**

File name: Supplementary_File_06_Reference_List

File format: .xls

Title of data: List of references included in the ontology

Description of data: References used to build up and annotate in the AO and the DO of *C. robusta* (*C. intestinalis* type A).

**Supplementary Data S7 is this one a supplementary data or supplementary table file?**

File name: Supplementary_File_07_Anatomical_Ontology

File format: .xls

Title of data: Anatomical Ontology of *C. robusta* (*C. intestinalis* type A) from Stage 26 to Stage 37

Description of data: Table listing the anatomical entities of the AO (Columns B-F: Terms and their specifications), their definition (Column G), Part of (Column H), Develops from (Column I), End Stage (Column J), Start Stage (Column K), Comment (Column L), Literature (Column M), and ID (Column N).

**Supplementary Data S8**

File name: Supplementary_File_08_Anatomical_Dictionary

File format: .xls

Title of data: List of anatomical entities, abbreviations, and definitions

Description of data: Table listing, in alphabetical order, the anatomical entities of the AO, their abbreviations used in Figures and Supplementary files, and their definitions.

**Supplementary Figure S9**

File name: Supplementary_File_09_Stage_26

File format: .jpg

Title of data: Hatching larva (Stage 26). CLSM

Description of data: **A.** Larva in the left lateral view. DCEN, RTEN, VCEN: cilia to dorsal caudal, rostral trunk, and ventral caudal epidermal neurons, respectively; pp: three anterior adhesive papillae. **B-B**^**I**^. Medial sagittal optic sections of the larval trunk (B) and posterior part of the tail (B^I^). Enlargement is the same in B and B^I^. **C-C**^**I**^. Transverse optic sections of larval trunk at sensory vesicle (C) and the atrial siphon primordia (C^I^) level. Enlargement is the same in C and C^I^. **D-D**^**I**^. Frontal optic sections of the larval trunk at the sensory vesicle (D) and ventral pharynx (D^I^) level. Enlargement is the same as C. ant pha: anterior pharynx; CNS: larval central nervous system; epi: epidermis; lasp: left atrial siphon primordium; mech: mesenchyme; ne: neck; nc: nerve cord; noto: notochord; oc: ocellus; osp: oral siphon primordium; noto: notochord; pha: pharynx; post pha: posterior pharynx; pp: papilla; rasp: right atrial siphon primordium; sv: sensory vesicle; vg: visceral ganglion.

**Supplementary Figure S10**

File name: Supplementary_File_10_Stage_28

File format: .jpg

Title of data: Mid swimming larva (Stage 28)

Description of data: **A.** Larva, left view. Note the ciliary network belonging to the epidermal sensory neurons (DCEN, RTEN, VCEN: cilium of a dorsal caudal, rostral trunk, and ventral caudal epidermal neuron, respectively). CLSM. **B.** Median sagittal optic section. CLSM. **C-C**^**II**^. Two transverse optic sections of the same larva of B. Enlargement is the same in D-D^I^. CNS: central nervous system; epi: epidermis; lasp: left oral siphon primordium; mech: mesenchyme; nc: nerve cord; nd: neurohypophyseal duct; ne: neck; noto: notochord; pl: preoral lobe;; osp: oral siphon primordium; pha: pharynx; pp: dorsal papillae; sv: sensory vesicle; vg: visceral ganglion.

**Supplementary Figure S11**

File name: Supplementary_File_11_Stage_29

File format: .jpg

Title of data: Late swimming larva (Stage 29)

Description of data: **A.** Larva, left view. CLSM. ATEN, DCEN, RTEN, VCEN: cilium of an anterior trunk, dorsal caudal, rostral trunk, and ventral caudal epidermal neuron, respectively. **B.** Medial sagittal optic section of the larval trunk. CLSM. **C-C**^**I**^. Two frontal sections of the same larva, at the level of atrial siphon primordia (C) and endodermal strand (esr) (C^I^). Light microscopy, Toluidine blue. Enlargement is the same in C-C^I^. **D-D**^**VII**^. Eight selected transverse sections from a complete dataset of a serially sectioned larva from anterior (D) to posterior (D^VII^). Light microscopy, Toluidine blue. Enlargement is the same in D-D^VII^. CNS: central nervous system; cor: coronet cells; epi: epidermis; esp: endostyle primordium; iclt (C2) and oclt (C1): inner (C2) and outer (C1) cuticular layer of the tunic, respectively; gp: gut primordium; lasp: left atrial siphon primordium; mech: mesenchyme; nc: nerve cord; noto: notochord; oc: ocellus; ot: otolith; pha lum: pharynx lumen; pl: preoral lobe; pp: ventral papilla; rasp: right atrial siphon primordium; sv: sensory vesicle; tc: test cell; tf: tail fin; tmc: tail muscle cells; tun: tunic; tunc: tunic cells; vg: visceral ganglion.

**Supplementary Figure S12**

File name: Supplementary_File_12_Stages_31-32.jpg

File format: .jpg

Title of data: Early tail absorption (Stage 31) (A-B^I^) and mid tail absorption (Stage 32) (C-F^II^). CLSM

Description of data: **A.** Larva in early tail absorption, seen from the left side. **B-B**^**I**^. Medial sagittal optic sections of the trunk (B) and tail tip (B^I^) of the larva shown in A. **C.** Larva in the mid-tail absorption, seen from the left side. The line on the larval trunk labeled by D^II^ indicates the level of the transverse section shown in D^II^. **D-D**^**I**^. Medial sagittal optic sections of the tail tip (D) and trunk (D^I^) of the larva shown in C. In D^I^, the line on the larval trunk labeled by D^II^ indicates the level of transverse section shown in D^II^. Enlargement is the same in D-D^I^. **D**^**II**^. Transverse optic section of the larval trunk shown in C and D^I^. **E-E^II^**. Larva in mid-tail absorption (E), seen from the left side, at a slightly more advanced stage than in C, and detail of its tail tip (E^I^, and its optic section in E^II^). The lines on the larval trunk labeled by F and F^II^ indicate the levels of sections shown in F and F^II^, respectively. Enlargement is the same in E-E^II^. **F-F**^**II**^. Transverse (F), medial sagittal (F^I^), and frontal (F^II^) optic sections of the trunk of larva shown in E. Enlargement is the same in F^I^-F^II^. abs tail: absorbing tail; ATEN, DCEN, RTEN, VCEN: cilium of an anterior trunk, dorsal caudal, rostral trunk, and ventral caudal epidermal neuron, respectively; deg pp: degenerating papilla; deg tail: degenerating tail; esr: endodermal strand; epi: epidermis; esp: endostyle primordium; gp: gut primordium; ht: heart; lasp: left atrial siphon primordium; lbr: larval brain remnants; mech: mesenchyme; oes: oesophagus; ne: neck; noto: notochord; pha lum: pharynx lumen; pl: preoral lobe; pp: ventral papilla; rasp: right atrial siphon primordium; sv: sensory vesicle; tf: tail fin; tmc: tail muscle cell; tunc: tunic cells; vg: visceral ganglion.

**Supplementary Figure S13**

File name: Supplementary_File_13_Stage_33.jpg

File format: .jpg

Title of data: Late tail absorption (Stage 33). CLSM

Description of data: **A-A**^**II**^. Larva in late tail adsorption (A), seen from the dorsal side, and optic frontal (A^I^) and transverse (A^II^) sections. The line on the larval trunk labeled by A^II^ indicates the level of the section shown in A^II^. Enlargement is the same in A-A^I^. **B-B**^**II**^. Larva in late tail absorption (B), seen from the dorsal side at a more advanced stage than in A, and optic frontal (B^I^) and transverse (B^II^) sections. The line on the larval trunk labeled by B^II^ indicates the level of the section shown in B^II^. Enlargement in B-B^I^ is the same as in A. abs tail: absorbing tail; brc: branchial chamber; esp: endostyle primordium; gp: gut primordium; gs: gill slit; hc: haemocyte; lasp: left atrial siphon primordium; os: oral siphon; rasp: right atrial siphon primordium.

**Supplementary Figure S14**

File name: Supplementary_File_14_Stage_35.jpg

File format: .jpg

Title of data: Mid body axis rotation (Stage 35)

Description of data: **A-A**^**IV**^. Metamorphosing larva, seen from the left side (A) and its medial sagittal (A^I^), frontal (A^II^), and transverse (A^III^) optic sections. A^IV^ is an enlargement of the squared area in A^I^. In A^I^, the lines on the larval trunk labeled by A^II^ and A^III^ indicate the levels of sections shown in A^II^ and A^III^, respectively. brc: branchial chamber; cil duc: ciliated duct of neural gland; es: endostyle; hc: haemocytes; ht: heart; las: left atrial siphon; lpsm: left protostigmata; ng: neural gland; os: oral siphon; osm: oral siphon muscle; psm: right protostigmata; stom: stomach; tail remn: tail remnants; tun: runic; tunc: tunic cells. Enlargement is the same in A and A^I^.

**Supplementary Figure S15**

File name: Supplementary_File_15_Stage_37.jpg

File format: .jpg

Title of data: Early juvenile I (Stage 37)

Description of data: **A-A**^**II**^. Juvenile, seen from the left side (A), its medial sagittal optic section (A^I^) and its depth-coded image. The depth information is represented by a heat map: warmer colors go to the front, and cooler colors to the back. For the color bar, see Fig. 5A^II^. Lines indicated by B-B^VI^ in A represent the level of transverse sections shown in B-B^VI^.CLSM. Enlargement is the same in A-A^II^. **B-B**^**VI**^. Transverse sections of a juvenile from the dorsal (B) to ventral (B^VI^) sides. Squared areas in B^III^ and B^IV^ are enlarged in insets to show details of endostyle (B^III^), left atrial siphon (black square), and row of ciliated cells of a protostigma (red square) (B^IV^), respectively. Numbers 1-8 in the inset of B^III^ indicate the eight zones of endostyle. Arrowheads in B^II^: ciliated cells of the coronal organ; asterisks in B^III^-B^IV^: protostigmata. Toluidine blue. Enlargement is the same in B-B^VI^. as: atrial siphon; brc: branchial chamber; cil duc: ciliated duct of neural gland; cg: cerebral ganglion; cut: tunic cuticle; dl: dorsal lamina; es: endostyle; hc: haemocyte; int: intestine; las: left atrial siphon; lasm: left atrial siphon muscles; lbr: larval brain remnant; man: mantle; mc: myocardium; mint: medium intestine; ng: neural gland; oes: oesophagus; os: oral siphon; osm: oral siphon muscle; pb: peripharyngeal band; pc: pericardium; pg: pyloric gland; prox int: proximal intestine; psm: protostigmata; pyc: pyloric caecum; stom: stomach; term int: terminal intestine, close to anus; ras: right atrial siphon; tail remn: tail remnants; ten: oral tentacles; tun: runic; tunc: tunic cells.

**Supplementary Table S16**

File name: Supplementary_File_16_available_ontologies.jpg

File format: .jpg

Title of data: List of anatomical ontologies

Description of data: Table of available ontologies regarding the anatomy of animal model organisms summarizing: the combination of AO with developmental stages, the number of anatomical terms listed, the percentage number of the relation “develops from” with respect to the total number of terms, the use of references as a source of data, and the link to the FAIRsharing database.

## References

1. Van Slyke, C. E., Bradford, Y. M., Westerfield, M. & Haendel, M. A. The zebrafish anatomy and stage ontologies: representing the anatomy and development of Danio rerio. J. Biomed. Semantics 5, 12 (2014).

2. Feric, M. et al. Coexisting Liquid Phases Underlie Nucleolar Subcompartments. Cell 165, 1686–1697 (2016).

3. Mungall, C. J., Torniai, C., Gkoutos, G. V., Lewis, S. E. & Haendel, M. A. Uberon, an integrative multi-species anatomy ontology. Genome Biol. 13, R5 (2012).

4. Manni, L. et al. Ontology for the asexual development and anatomy of the colonial chordate botryllus schlosseri. PLoS One 9, (2014).

5. Segerdell, E. et al. Enhanced XAO: the ontology of Xenopus anatomy and development underpins more accurate annotation of gene expression and queries on Xenbase. J. Biomed. Semantics 4, 31 (2013).

6. Bourlat, S. J. et al. Deuterostome phylogeny reveals monophyletic chordates and the new phylum Xenoturbellida. Nature (2006). doi:10.1038/nature05241

7. Delsuc, F., Brinkmann, H., Chourrout, D. & Philippe, H. Tunicates and not cephalochordates are the closest living relatives of vertebrates. Nature (2006). doi:10.1038/nature04336

8. Satoh, N. The ascidian tadpole larva: Comparative molecular development and genomics. Nat. Rev. Genet. 4, 285–295 (2003).

9. Corbo, J. C., Di Gregorio, A. & Levine, M. The Ascidian as a Model Organism in Developmental and Evolutionary Biology. Cell 106, 535–538 (2001).

10. Procaccini, G., Affinito, O., Toscano, F. & Sordino, P. A New Animal Model for Merging Ecology and Evolution. in Evolutionary Biology – Concepts, Biodiversity, Macroevolution and Genome Evolution 91–106 (Springer Berlin Heidelberg, 2011). doi:10.1007/978-3-642-20763-1_6

11. Gallo, A. & Tosti, E. The Ascidian Ciona Intestinalis as Model Organism for Ecotoxicological Bioassays. J. Mar. Sci. Res. Dev. (2015). doi:10.4172/2155-9910.1000e138

12. Caputi, L. et al. Cryptic speciation in a model invertebrate chordate. Proc. Natl. Acad. Sci. U. S. A. (2007). doi:10.1073/pnas.0610158104

13. Nydam, M. L. & Harrison, R. G. Genealogical relationships within and among shallow-water Ciona species (Ascidiacea). Mar. Biol. (2007). doi:10.1007/s00227-007-0617-0

14. Nydam, M. L. & Harrison, R. G. Polymorphism and divergence within the ascidian genus Ciona. Mol. Phylogenet. Evol. (2010). doi:10.1016/j.ympev.2010.03.042

15. Nydam, M. L. & Harrison, R. G. Introgression despite substantial divergence in a broadcast spawning marine invertebrate. Evolution (N. Y). (2011). doi:10.1111/j.1558-5646.2010.01153.x

16. Nydam, M. L. & Harrison, R. G. Reproductive protein evolution in two cryptic species of marine chordate. BMC Evol. Biol. (2011). doi:10.1186/1471-2148-11-18

17. Sato, A., Satoh, N. & Bishop, J. D. D. Field identification of ‘types’ A and B of the ascidian Ciona intestinalis in a region of sympatry. Marine Biology (2012). doi:10.1007/s00227-012-1898-5

18. Sato, A., Shimeld, S. M. & Bishop, J. D. D. Symmetrical reproductive compatibility of two species in the ciona intestinalis (ascidiacea) species complex, a model for marine genomics and developmental biology. Zoolog. Sci. (2014). doi:10.2108/zs130249

19. Roux, C., Tsagkogeorga, G., Bierne, N. & Galtier, N. Crossing the species barrier: Genomic hotspots of introgression between two highly divergent ciona intestinalis species. Mol. Biol. Evol. (2013). doi:10.1093/molbev/mst066

20. Bouchemousse, S., Bishop, J. D. D. & Viard, F. Contrasting global genetic patterns in two biologically similar, widespread and invasive Ciona species (Tunicata, Ascidiacea). Sci. Rep. 6, 1–15 (2016).

21. Brunetti, R. et al. Morphological evidence that the molecularly determined Ciona intestinalis type A and type B are different species: Ciona robusta and Ciona intestinalis. J. Zool. Syst. Evol. Res. 53, 186–193 (2015).

22. Pennati, R. et al. Morphological Differences between Larvae of the Ciona intestinalis Species Complex: Hints for a Valid Taxonomic Definition of Distinct Species. PLoS One 10, e0122879 (2015).

23. Gissi, C. et al. An unprecedented taxonomic revision of a model organism: the paradigmatic case of Ciona robusta and Ciona intestinalis. Zoologica Scripta (2017). doi:10.1111/zsc.12233

24. Hotta, K. et al. A web-based interactive developmental table for the Ascidian Ciona intestinalis, including 3D real-image embryo reconstructions: I. From fertilized egg to hatching larva. Dev. Dyn. 236, 1790–1805 (2007).

25. Whetzel, P. L. et al. BioPortal: Enhanced functionality via new Web services from the National Center for Biomedical Ontology to access and use ontologies in software applications. Nucleic Acids Res. (2011). doi:10.1093/nar/gkr469

26. Conklin, E. G. Mosaic development in ascidian eggs. J. Exp. Zool. 2, 145–223 (1905).

27. Nishida, H. Cell Lineage Analysis in Ascidian Embryos by Intracellular Injection of a Tracer Enzyme III. Up to the Tissue Restricted Stage. Dev. Biol. 121, 526–541 (1987).

28. Nishida, H. & Satoh, N. Cell Lineage Analysis in Ascidian Embryos by Intracellular Injection of a Tracer Enzyme I. Up to the Eight-Cell Stage. Dev. Biol. 99, 382–394 (1983).

29. Satoh, N. CELLULAR MORPHOLOGY AND ARCHITECTURE DURING EARLY MORPHOGENESIS OF THE ASCIDIAN EGGLJ: AN SEM STUDY. Biol. Bull. 155, 608–614 (1978).

30. Cloney, R. A. Ascidian Larvae and the Events of Metamorphosis. Am. Zool. 22, 817–826 (1982).

31. Willey, A. Memoirs: Studies on the Protochordata. Quart. J. Micr. Sci. s2-34, 317–360 (1893).

32. R. H. Millar. XXXV. CIONA. in L.M.B.C 1–123 (1953).

33. Chiba, S., Sasaki, A., Nakayama, A., Takamura, K. & Satoh, N. Development of Ciona intestinalis juveniles (through 2nd ascidian stage). Zoolog. Sci. 21, 285–298 (2004).

34. Day-Richter, J., Harris, M. A., Haendel, M. & Lewis, S. OBO-Edit an ontology editor for biologists. Bioinformatics 23, 2198–2200 (2007).

35. Nishida, H. Cell lineage analysis by intracellular injection of a tracer enzyme. Dev. Biol. 121, 526–541 (1987).

36. Taniguchi, K. & Nishida, H. Tracing cell fate in brain formation during embryogenesis of the ascidian Halocynthia roretzi. Dev. Growth Differ. 46, 163–180 (2004).

37. Hudson, C. The central nervous system of ascidian larvae. Wiley Interdiscip. Rev. Dev. Biol. 5, 538–561 (2016).

38. Wakai, M. K., Nakamura, M. J., Sawai, S., Hotta, K. & Oka, K. Two-Round Ca2+ Transients in Papillae by Mechanical Stimulation Induces Metamorphosis in the Ascidian,. submitted (2020).

39. Matsunobu, S. & Sasakura, Y. Time course for tail regression during metamorphosis of the ascidian Ciona intestinalis. Dev. Biol. 405, 71–81 (2015).

40. Richardson, L. & Armit, C. Digital Graphical Resources and Developmental Anatomy in the Mouse. in Kaufman’s Atlas of Mouse Development Supplement (2016). doi:10.1016/b978-0-12-800043-4.00024-5

41. Burighel, P., Cloney, R. A. & Cloney, B. Microscopic Anatomy of Invertebrates, Vol. 15. Microsc. Anat. Invertebr. 15, 221–347 (1997).

42. Chambon, J.-P. P. et al. Tail regression in Ciona intestinalis (Prochordate) involves a Caspase-dependent apoptosis event associated with ERK activation. Development 129, 3105–3114 (2002).

43. Kott, P. The Australian Ascidiacea. Part 1, Phlebobranchia and Stolidobranchia. Mem. Queensl. Museum. (1985).

44. Hirano, T. & Nishida, H. Developmental Fates of Larval Tissues after Metamorphosis in AscidianHalocynthia roretzi. Dev. Biol. 192, 199–210 (1997).

45. Stolfi, A. et al. Divergent mechanisms regulate conserved cardiopharyngeal development and gene expression in distantly related ascidians. Elife 3, e03728 (2014).

46. Wang, W., Razy-Krajka, F., Siu, E., Ketcham, A. & Christiaen, L. NK4 Antagonizes Tbx1/10 to Promote Cardiac versus Pharyngeal Muscle Fate in the Ascidian Second Heart Field. PLoS Biol. 11, e1001725 (2013).

47. Hirano, T. & Nishida, H. Developmental fates of larval tissues after metamorphosis in the ascidian, Halocynthia roretzi. II. Origin of endodermal tissues of the juvenile. Dev. Genes Evol. 210, 55–63 (2000).

48. Ohtsuka, Y., Matsumoto, J., Katsuyama, Y. & Okamura, Y. Nodal signaling regulates specification of ascidian peripheral neurons through control of the BMP signal. Development 141, 3889–3899 (2014).

49. Sansone, S.-A. et al. FAIRsharing as a community approach to standards, repositories and policies. Nat. Biotechnol. 37, 358–367 (2019).

50. Essock-Burns, T. et al. Barnacle biology before, during and after settlement and metamorphosis: a study of the interface. J. Exp. Biol. 220, 194–207 (2017).

51. Kitamura, H., Kitahara, S. & Koh, H. B. The induction of larval settlement and metamorphosis of two sea urchins, Pseudocentrotus depressus and Anthocidaris crassispina, by free fatty acids extracted from the coralline red alga Corallina pilulifera. Mar. Biol. 115, 387–392 (1993).

