## Supplementary figures and images for "The ontology of the anatomy and development of the solitary ascidian *Ciona*"

### Supplementary_Figure_01_Life_Cycle

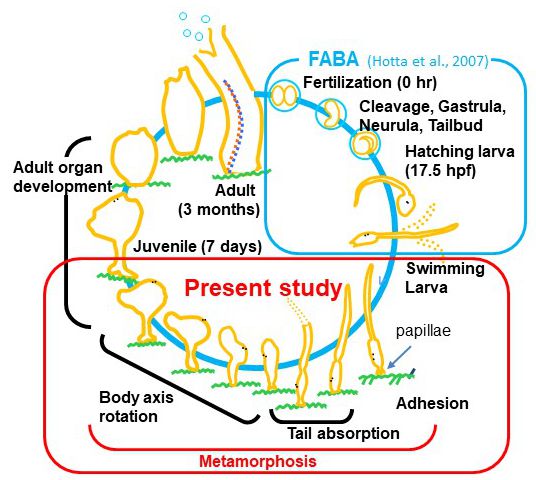

### Supplementary_File_05_3D

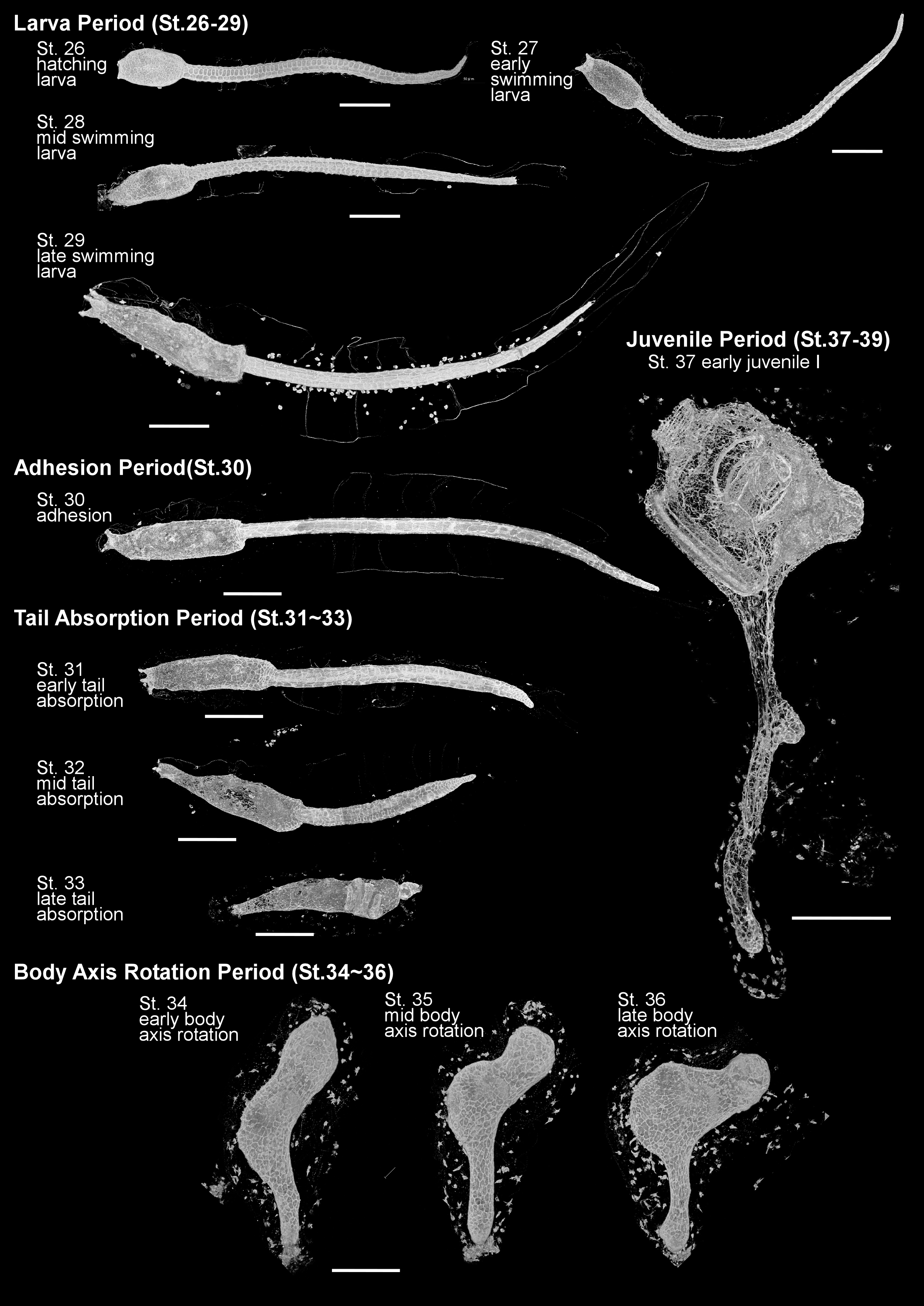

### Supplementary_File_09_Stage_26

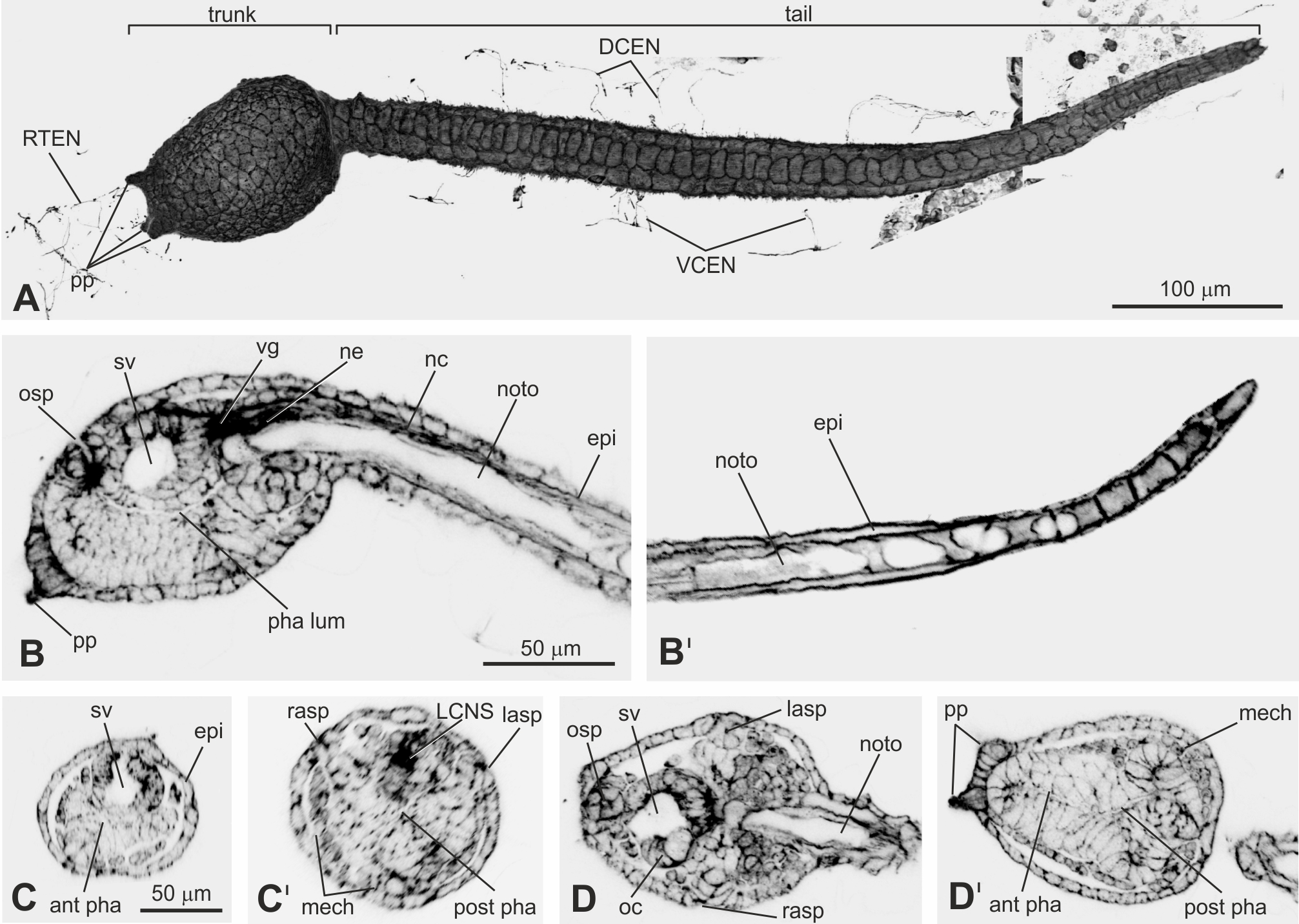

### Supplementary_File_10_Stage_28

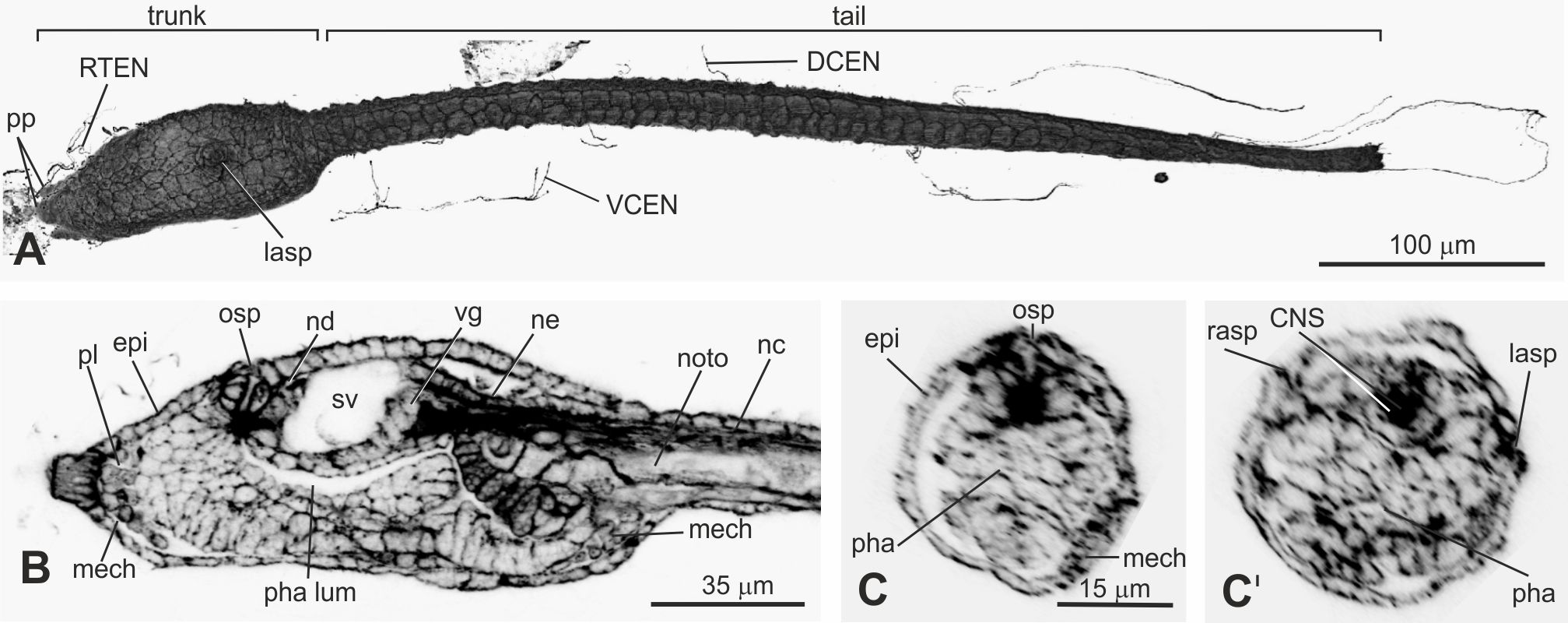

### Supplementary_File_11_Stage_29

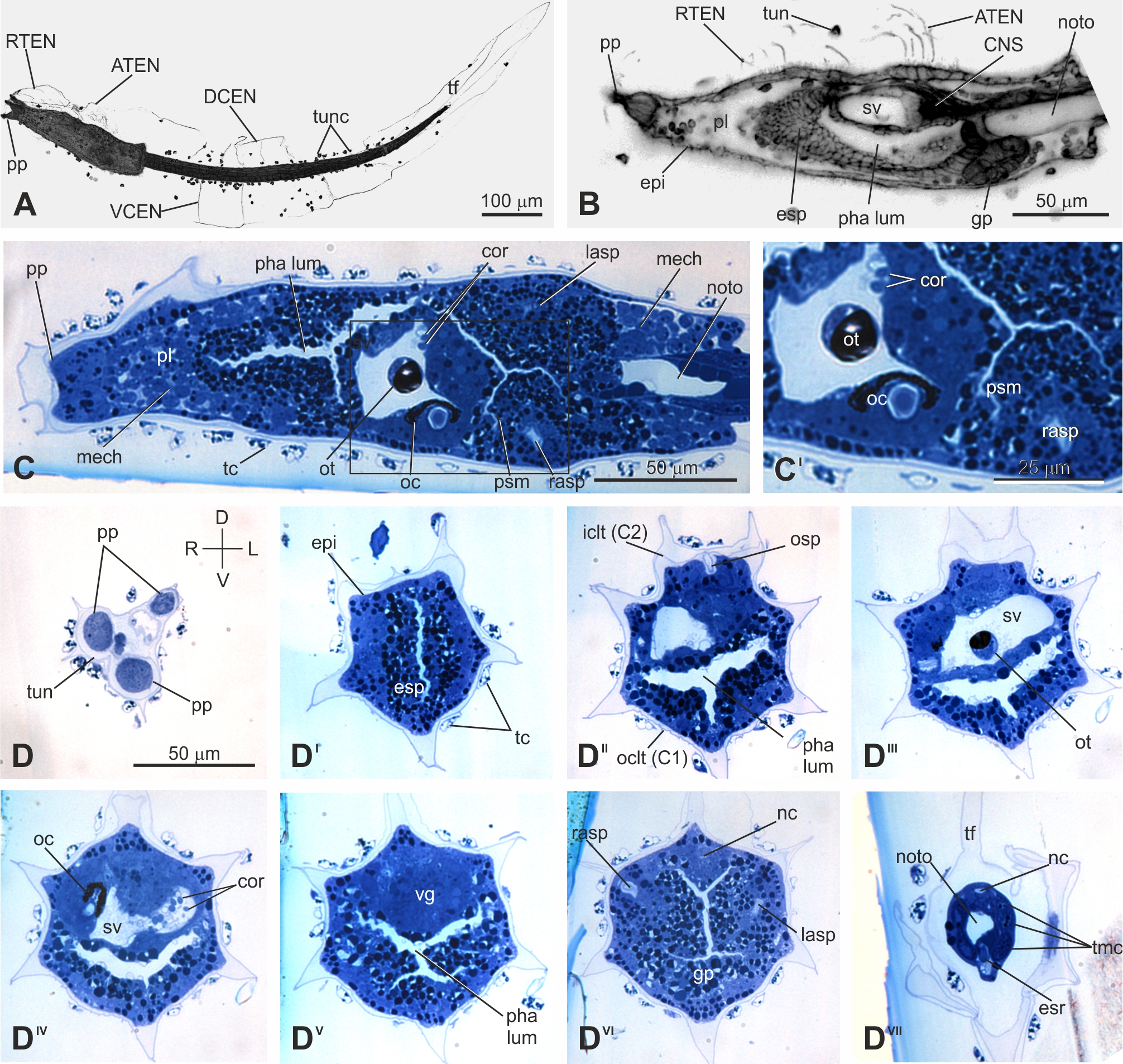

### Supplementary_File_12_Stages_31-32

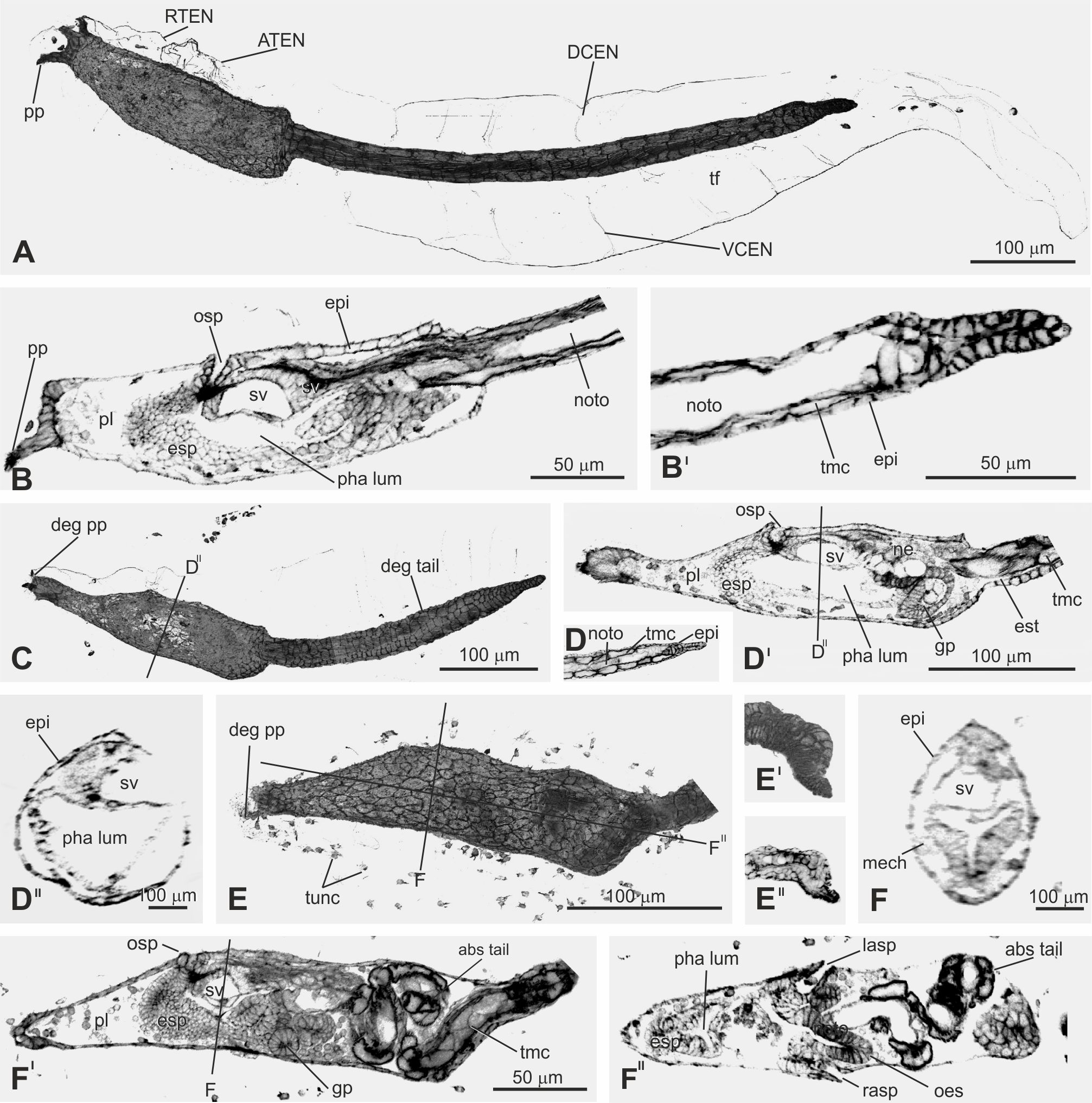

### Supplementary_File_13_Stage_33

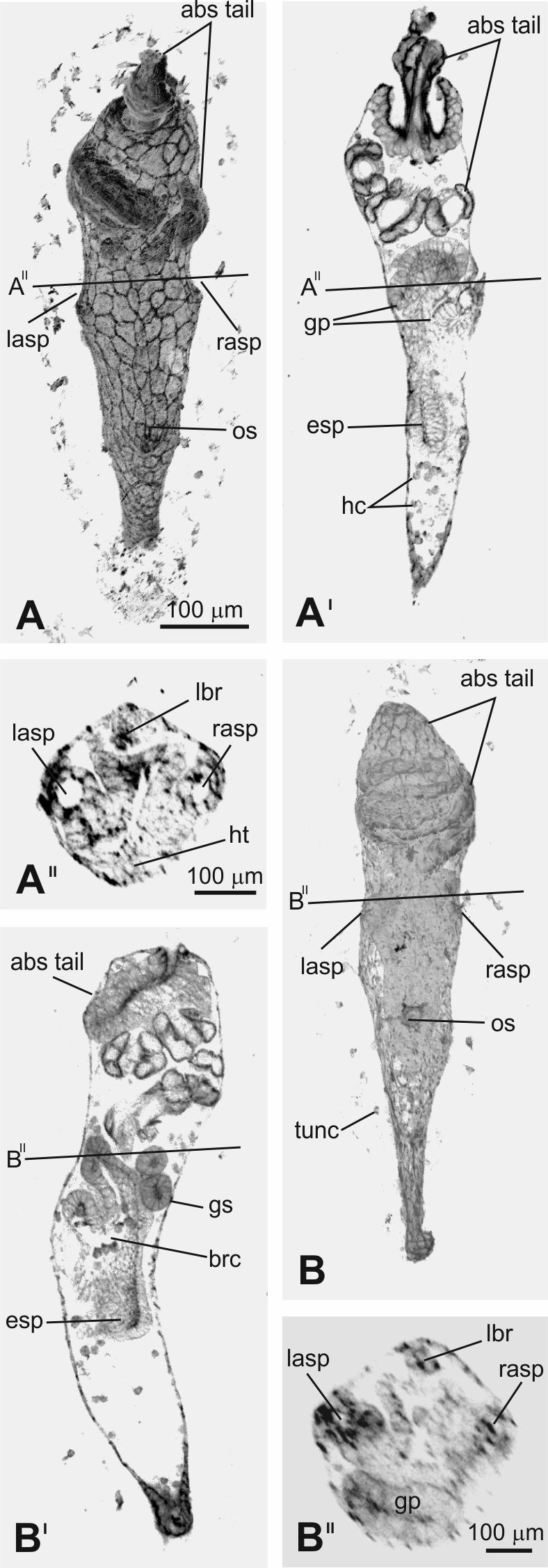

### Supplementary_File_14_Stage_35

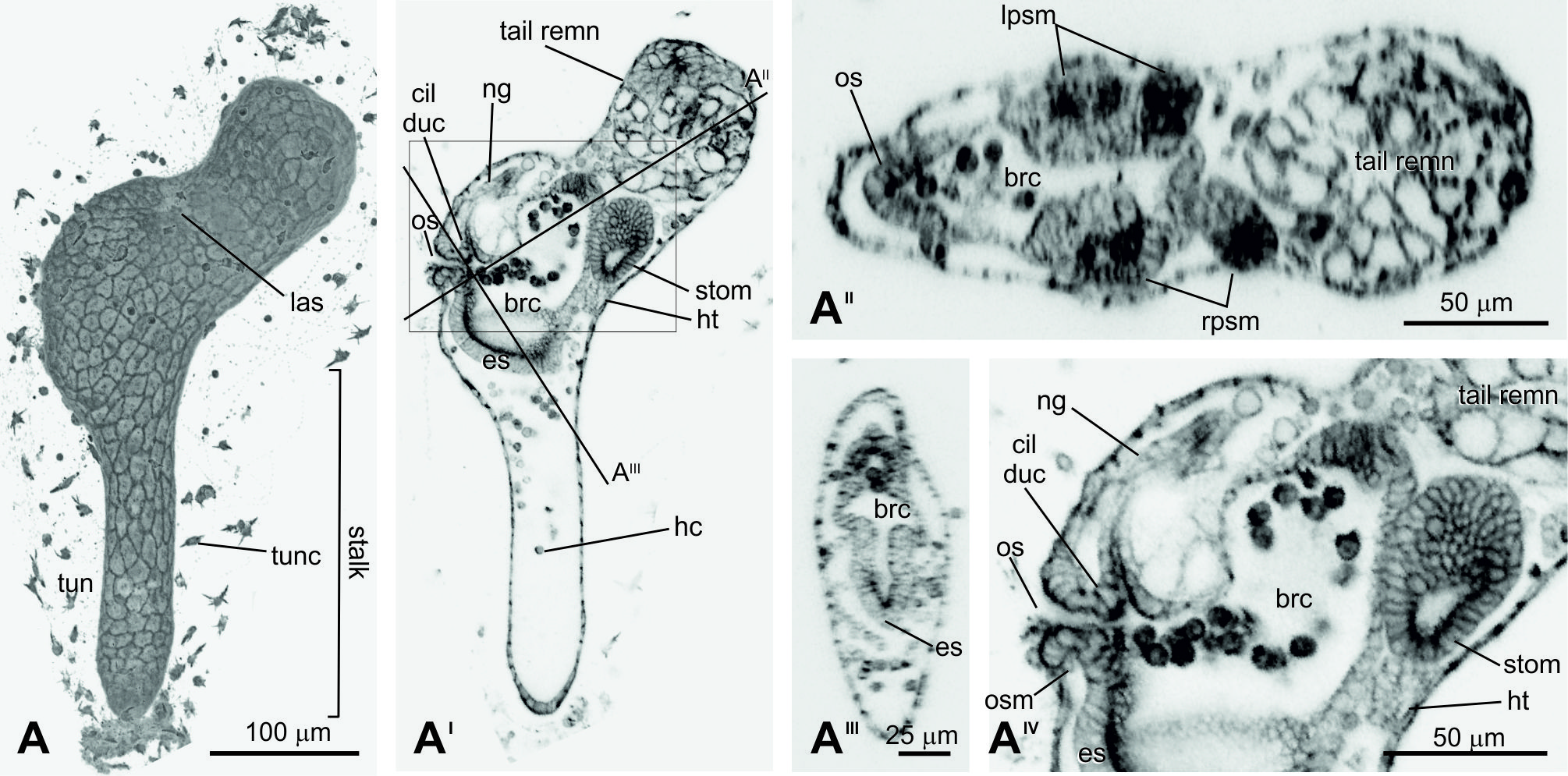

### Supplementary_File_15_Stage_37

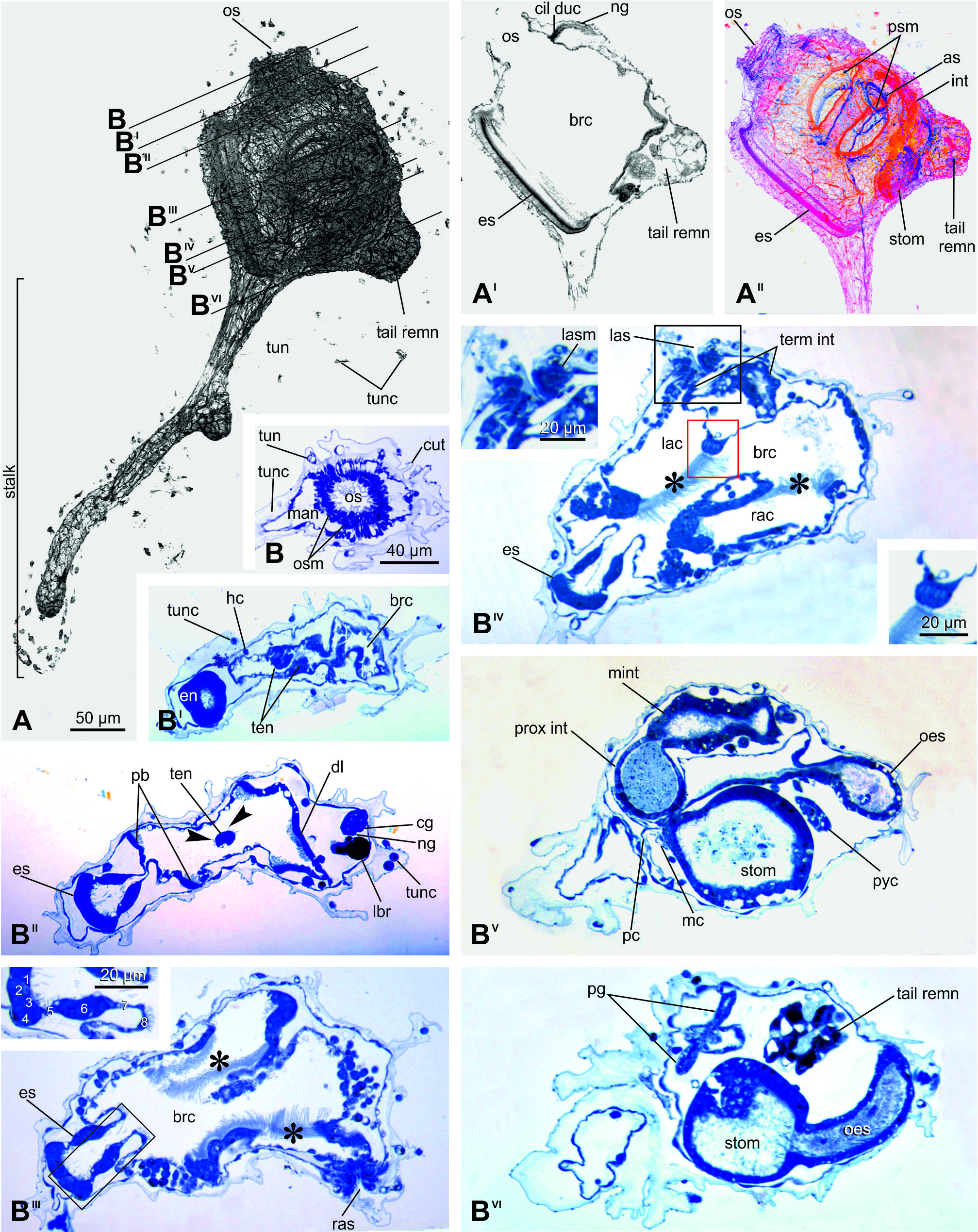

### Supplementary_File_16_available_ontologies

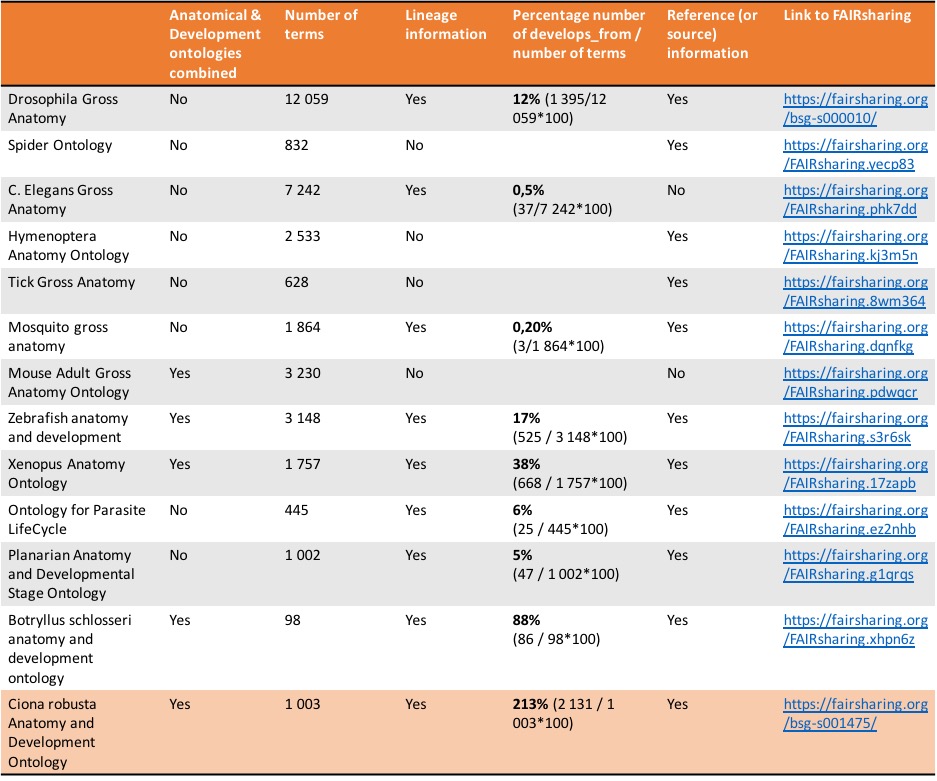
