## Supplementary_File_02_DescriptionStages2 for "The ontology of the anatomy and development of the solitary ascidian *Ciona*"

Supplementary Data S2

Stage 26

This is the **Hatching Larva Stage** (CirobuD:0000049; 17.5 h, Supplementary Figure S9). The larva has a roundish trunk and immature papillae (*pp* in S9A, B, D^I^; CirobuA:0000675) with round tips; it exhibits irregular **tail** (CirobuA:0000797) movements. Eight strips of **epidermal cells** (*epi* in S9B, B’, C; CirobuA:0000594) constitute the **tail epidermis** (CirobuA:0000738). The larval **pharynx** (*pha* in S9B; CirobuA:0000682) shows a narrow lumen ^1^. The larval nervous system is subdivided into a central and peripheral nervous system. The former, as described above, contains several entities, for example, the otolith and the ocellus with **lens cells** (CirobuA:0000648), **photoreceptors** (CirobuA:0000683), and **pigment cup cells** (CirobuA:0000685); and the visceral ganglion (*vg* in S9B) with its motor neurons. The **larval peripheral nervous system** (CirobuA:0000681) is differentiating and each epidermal sensory neuron, *i.e.*, the **Rostral Tail Epidermal Neurons** (*RTEN* in S9A; CirobuA:0000709), the **Dorsal Caudal Epidermal Neurons** (*DCEN* in S9A; CirobuA:0000589), and the **Ventral Caudal Epidermal Neurons** (*VCEN* in S9A; CirobuA:0000757), shows dendritic arbors. A vacuolated notochord (*noto* in S9B, B^I^, D) extends as an elongated structure in the tail. The **oral siphon primordium** (*osp* in S9B; CirobuA:0000670) is in the form of an epidermal invagination not yet communicating with the pharynx lumen. The **atrial siphon primordia** (*rasp* and *lasp* in S9C^I^; CirobuA:0000368), the pair of invaginations of the dorsal-lateral trunk epidermis, called also **atrial placodes** (CirobuA:0000908), are not yet open to the exterior. The trunk is covered by a double-layered **tunic** (CirobuA:0000750) that, at this stage, is not well distinguishable at the histological level: the inner compartment of the tunic (CirobuA:0000882), bordered by the **inner cuticular layer** (CirobuA:0000883), and the **outer compartment of the tunic** (CirobuA:0000884), bordered by the **outer cuticular layer** (CirobuA:0000885).

Stage 27

This corresponds to the **Early Swimming Larva** **Stage** (CirobuD:0000050; 17.5-20 h; Fig. 2). The larva makes regular tail movements while swimming. The trunk elongates along the anterior-posterior axis. The **postpharyngeal tract** (*post pha* in Fig. 2C^II^; CirobuA:0000871) is developing from the posterior pharynx and the **endodermal strand** ^2^ (*est* in Fig. 2C^III^) is histologically recognizable.

The **endostyle primordium** (*est* in C^III^; CirobuA:0000617), *developing from* A7.1, A7.2, A7.5, B7.1, and B7.2 cells ^1^, is recognizable in the anterior pharynx (*ant pha* in Fig. 2B^II^). A wider lumen of the pharynx can be observed (compare *pha lum* in B of Supplementary Figure S9 and Fig. 2C^II^ E^I^). The atrial siphon primordia indent in the **posterior lateral trunk** **epidermis** (a7.14 and a7.15 cell lines) (CirobuA:0000916; compare *last* and *rasp* in C^I^ in Supplementary Figure S9 with Fig. 2D^II^, E, E^II^). Mesenchyme cells (*mech* in Fig. 2B^I^, C^I^, C^II^, D) in the posterior ventral trunk become round. Among sensory structures in the sensory vesicle (*sv* in B^I^, C, C^I^, C^II^), the **coronet cells** (*cor* in Fig. 2B^II^; CirobuA:0000890) can also be noted. These cells, forming a hydropressure organ ^3,4^, are considered by some authors to be the homolog of the vertebrate hypothalamus ^5–8^.

Stage 28

This is the **Mid-Swimming Larva** **Stage** (CirobuD:0000051; 20-22 h; Supplementary Figure S10). The papillae (*pp* in S10A-C^II^) elongate and their basal part expands. The trunk is square in shape. The ciliary network belonging to the epidermal sensory neurons (**ascidian dendritic network in tunic**, or ASNET; CirobuA:0000892) becomes more complex (*ATEN*, *DCEN*; *RTEN*, and *VCEN* in S10A). The **preoral lobe** (CirobuA:0000901; *pl* in S10B), a wide anterior body cavity between the pharynx and the anterior epidermis, is recognizable. Here, round mesenchyme cells are present (*mech* in S10B). Spaces among the epithelia (**haemocele**, CirobuA:0000888) become larger; here, mesenchyme cells represent the **haemocyte** (CirobuA:0000571; synonym of blood cell) precursors. According to Parrinello and co-authors ^8^, haemocytes in adult animals are stem cells and granulocytes. The latter include clear granulocytes (precursors to clear vesicular granulocytes), microgranulocytes, and vacuolar granulocytes (including unilocular granulocytes and globular granulocytes). **Tunic cells** (CirobuA:0000751) are also recognizable in the inner compartment of the tunic. They differentiate from mesenchyme cells. Coronet cells are well-differentiated on the left side of the sensory vesicle. The **otolith** (CirobuA:0000671) senses gravity and gravitaxis is observed at this stage ^9^.

Stage 29

This is the **Late Swimming Larva** **Stage** (CirobuD:0000052; 22-24 h; Supplementary Figure S11). With respect to the previous stage, the trunk is longer and narrower, and the tail is longer. Moreover, the trunk profile is squared at the trunk-tail transition (S11A, C-C^I^); in cross-section, the trunk and its tunic (*tun* in S11D) are polygonal and star-shaped, respectively (see S11 D-D^vii^). All of the larval structures for swimming are fully mature: the **tunic fin** (*tf* in S11A, D^IIV^; CirobuA:0000622) located along the dorso-ventral axis, the **tail muscle** (*tmc* in S11D^IIV^; CirobuA:0000739) fibers, and the tunic ciliated sensory fields (*ATEN*, *DCEN*; *RTEN*, and *VCEN* in S11A-B;Terakubo et al., 2010; Yokoyama et al., 2014). In *Ciona*, the larva swims by tail locomotion for several hours. The gravitaxis and visual behaviors are tightly interconnected at this stage ^9^. The duration of the swimming period is variable among individuals. In our observations, it lasts until 23.6 hpf on average, *i.e.*, until the adhesion at the beginning of metamorphosis.

The **gut primordium** (synonym: “gut rudiment”; *gp* in S11D^VI^; CirobuA:0000862) is histologically recognizable as endodermal tissue posterior to the pharynx. Here, the **protostigmata** (CirobuA:0000870) rudiments are now recognizable (*psm* in S11C; ^2^). According to Hirano and Nishida (2000), in *Halocynthia roretzi*, the atrial epithelium *develops from* A7.2, A7.1, A7.5, and B7.1 cell lines. It gives rise to the paired **atrial siphons** (CirobuA:0000366), not yet in communication with the **atrial cavities** (CirobuA:0000798) (*lasp* and *rasp* in S11C, D^VI^).

Stage 30

The **Adhesion Stage** (CirobuD:0000053) regards the larva attaching to a suitable substrate through its adhesive papillae (24-27 h; Fig. 3). The papillae change significantly after adhesion (Fig. 3A-C), becoming deflated and sometimes curved. They also start to degenerate. At this stage, the adhesion area is flat. Also called **holdfast** ^1^, in the successive stages it will elongate in the **stalk** (CirobuA:0000632), the epidermal peduncle by which the juvenile is attached to the substratum. The stalk possesses a cavity, derived from the preoral lobe (*pl* in Fig. 3B, B^I^, C^I^), with its own **blood** (CirobuA:0000571) circulation.

The two atrial siphons are now in continuity with the atrial cavities. In the juvenile (Stage 41), the siphons will fuse, becoming a single, dorsal atrial siphon. The sensory organs are recognizable within the sensory vesicle but are beginning to degenerate (*ot* and *oc* in C^III^ and C^vi^).

The inner and outer compartment of the tunic, with their inner and outer cuticular layer, respectively, are well recognizable (*ict* and *oct*, *iclt1*(*C1)* and *oclt*(*C2*), respectively, in Fig. 3C-C^I^).

Stage 31

At the **Early Tail Absorption Stage** (CirobuD:0000054) 27-8 hpf; Supplementary Figure S12 A-B^I^), the shrinkage of the **tail epidermis** (*epi* in S12B^I^; CirobuA:0000738) begins at the tail tip (**Tail tip epidermis**: CirobuA:0000949) (compare S12B^I^with S9B^I^). In the same time, the tail epidermal cells, originally flat, change into thick and cuboidal (*epi* in S12B^I^). The actin staining in the posterior tail is relatively strong, indicating the actin’s involvement in the shrinkage process ^10^. In addition, the tail inner tissues, such as the notochord (*noto* in S12B), the endodermal strand (*est* in S12D^I^), and the muscles (*tmc* in S12D^I^), begin to be arranged irregularly in the posterior tail region. In some individuals, the tail slightly bends at the trunk-tail transition (Fig. 1B, Stage 31). The otolith and ocellus are still present, although the larval brain is degenerating; their remnants will be recognizable during the Juvenile Period. The papillae, with their **dorsal palp neurons** (a8.18 line; CirobuA:0000601) and **ventral palp neurons** (a8.20 line; CirobuA:0000771), are no longer histologically recognizable at the end of the stage (*End stage*).

Stage 32

This corresponds to the **Mid Tail Absorption Stage** (CirobuD:0000055) 28-29 h; Supplementary Figure S12C-F^III^), when 50% of the tail has been resorbed into the larval trunk. The tail is shorter and thicker than in the previous stage (*deg tail* in S12C). The tail muscles (*tmc* in S12D^I^, F^I^) contract together with the notochord; they fold and stack into the posterior trunk (, *abs tail* in S12F^I-II^). On the other hand, the **tail epidermis** (CirobuA:0000738) contracts without folding and finally invaginates into the trunk region. The process recalls the one described during the absorbing of the tail of *Halocynthia roretzi* and *Botryllus schlosseri* ^11,12^. Although the contribution of apoptosis is not excluded ^13^, it has been suggested that the tail epidermis and the extracellular **notochord sheath** (CirobuA:0000946) generate the strong forces retracting the axial organs into the trunk ^12^. In fact, the tail absorption is inhibited by cytochalasin B, indicating that actin fibers play an important role in the process ^14^.

Concomitantly with tail absorption, the ASNET lose their organization ^15^.

The gut continues its differentiation. The **oral siphon** (*osp* in S12D^I^, F^I^; CirobuA:0000668) opens. The **oesophagus** (*oes* in S12F^II^; CirobuA:0000621) ^2,16^, the **stomach** (CirobuA:0000737) ^1,2,17^, and the **intestine** (CirobuA:0000635) ^2^ are histologically well recognizable. Also, the **heart** (CirobuA:0000628) is visible ^18–22^.

The **test cells** (CirobuA:0000915) are no longer present. They were originally encased in superficial depressions of the developing oocyte by the vitelline coat ^17,22–24^. After fertilization, they were moved into the perivitelline space, to attach to the outer cuticular layer of the tunic. At Stage 32, they are eliminated together with the outer tunic compartment layer and the outer cuticular layer. The stalk starts to elongate.

Stage 33

This corresponds to the **Late Tail Absorption Stage** (CirobuD:0000056; 29-30 h; Supplementary Figure S13), during which the tail becomes completely absorbed. Together with the tail, the 88 larval entities associated with the embryonic and larval stages are no longer recognizable (Fig. 6).

Both the notochord and the tail muscles are folded several times and coiled into the posterior trunk (*abs tail* in S13A-A^I^, B-B^I^). Moreover, the posterior trunk epidermis wraps around the absorbing tail. Strong actin staining can still be observed in the absorbing axial organ (muscles and notochord) and in the degenerating tail epidermis.

In histological sections, several newly formed structures associated with the juvenile lifestyle can be defined. Some of them are the **body wal**l (CirobuA:0000857), the **atrial siphon muscles** (CirobuA:0000367), and the **pericardial cavity** (CirobuA:0000678) ^16,20,21,25^. The stalk continues to elongate. Tunic cells *(tunc* in S13B) are numerous in the definitive tunic (the original inner compartment of the tunic with its inner cuticular layer).

Stage 34

At the **Early Body Axis Rotation Stage** (CirobuD:0000057; 30-36 hpf; Fig. 4), the stalk (Fig. 4A, B^III^) continues to elongate and forms, with the endostyle axis, an angle of about 90° (Fig. 1B, Stage 34). The strong actin intensity in the trunk region associated to the tail remnants (*tail remn* in Fig. 4A^I^, A^IV^) indicates that the latter is tightly packed.

Adult organs proceed with their differentiation. In the digestive system, the **pyloric caecum** (*pyc* in Fig. 4B^II^; CirobuA:0000630) is differentiating evagination of the stomach (*stom* in Fig. 4A^II^, B^III^-B^IV^). The latter starts to enlarge.

The **oral cavity** (CirobuA:0000802), representing the **oral siphon lumen** (*os* in A^III^, B-B^I^), extends to the rim of the **velum** (*ve* in Fig. 4B; CirobuA:0000900) and **tentacles** (CirobuA:0000741, now recognizable ^26–29^. The oral siphon lumen is in continuity with the **branchial chamber** (*brc* in Fig. 4A^II-IV^, B^I-II^; CirobuA:0000863; synonym: branchial sac, branchial cavity) lumen. Tunic cells are also in the tunic covering the inner oral siphon epidermis.

Stage 35

At the **Mid Body Axis Rotation Stage** (CirobuD:0000058; 6-45 h; Supplementary Figure S14), the endostyle axis is perpendicular to the axis passing through the stalk (Fig.1B, Stage 35). In the branchial chamber (*brc* in S14A^I-IV^), which is more expanded than in the previous stage, one pair of elliptical **gill-slits** (*lpsm* and *rpsm* in S14A^II^; CirobuA:0000623), separated by a **transverse bar** (CirobuA:0000746), allows for filtration. The transverse bar contains the transverse sinus of the branchial sac. The **peripharyngeal band** (CirobuA:0000680), which is the ciliated band of the pharynx delimiting the **prebranchial zone** (CirobuA:0000869) from the branchial one, is visible ^30^.

Stage 36

At the **Late Body Axis Rotation Stage** (CirobuD:0000059) 45-60 h (2 dpf); Fig. 5), the angle between the endostyle axis and the axis passing through the stalk is 30°- 60° (Fig. 1B, Stage 36). With respect to the previous stage, some new entities are now recognizable. At the gut level, the **pyloric gland** (*pg* in Fig. 5D^I^; CirobuA:0000705) emerged from the pyloric caecum. At the neural system level, the **neural complex** (CirobuA:0000659), composed of the **cerebral ganglion** (CirobuA:0000582), and the **neural gland complex** (CirobuA:0000661) are distinguishable. The latter is formed of the **neural gland body** (*ng* in Fig. 5B, D), which anteriorly exhibits a gland aperture, the **ciliated funnel** (CirobuA:0000584). The latter is located in the **dorsal tubercle** (CirobuA:0000932), on the roof of the prebranchial zone. Posteriorly, the neural gland body elongates into the **dorsal strand** (*dst* in Fig. 5D; CirobuA:0000930). In the adult, a **dorsal strand plexus** (CirobuA:0000931) extends along the dorsal strand. Some **nerves** (CirobuA:0000929) are elongating from the neurons located in the cerebral ganglion.

The filter-feeding activity starts at this stage. Consequently, multiple entities linked to the respiratory and alimentary tract become physiologically functioning. Food, brought by water entering the oral siphon, reaches the branchial cavity (*brc* in Fig. 5A^I^, B-C, D^I^) (delimited by the **branchial epithelium** (CirobuA:0000677)) and passes through the oesophagus (*oes* in Fig. 5D^I^), the stomach (*stom* in Fig. 5A^I-II^, B-B^I^, D^II^) and the intestine (divided into the **proximal** (*prox int* in Fig. 5D^II^; CirobuA:0000872), **mid** (*mint* in Fig. 5D^I^; CirobuA:0000655), and **distal intestine** (CirobuA:0000631)) for digestion. Fecal pellets are eliminated through the **anus** (CirobuA:0000362), which opens into the atrial chamber. In the branchial chamber, the endostyle (*es* in Fig.5A^I-II^, B-B^I^, D, D^III^) is now characterized by its zones in the form of eight symmetrical longitudinal cellular bands (from the median **zone 1** (CirobuA:0000782) to peripheral **zone 8** (CirobuA:0000790))^1,22^. It is involved in mucus production for filtration. The oral mechanoreceptor, the **coronal organ** (CirobuA:0000923), is developing on the oral tentacles and the velum. It controls the circulating seawater inside the animal body, together with the atrial **cupular organ** (CirobuA:0000924). The **circular muscular system** (CirobuA:0000859) and **longitudinal muscular system** (CirobuA:0000860), responsible for body contraction, become recognizable in the body wall. The heart (*ht* in Fig. 5A^I^, B-B^I^, D^III^), with its inner contractile **myocardium** (*mc* in Fig. 5D^III^; CirobuA:0000886) and outer **pericardium** (*pc* in Fig. 5D^III^; CirobuA:0000679) joined by a **rafe** (*rph* in Fig. 5D^III^; CirobuA:0000887), is now beating.

Stage 37

This is the **Early Juvenile I Stage** (CirobuD:0000060; Supplementary Figure S15), which occurs when the endostyle axis is almost parallel to the axis passing through the stalk (63-72 h (3 dpf); Fig. 1B, Stage 37). The stomach swells and the larval tail remnants are no longer present 4 dpf.

The hermaphrodite **reproductive system** (CirobuA:0000909) is now recognizable. The **female reproductive system** (CirobuA:0000910) is formed by a sac-like **ovary** (CirobuA:0000672) continuous in an **oviduct** (CirobuA:0000801). The **male reproductive system** (CirobuA:0000910) comprises the lobular **testis** encrusting the ovary (CirobuA:0000742) and the **sperm duct** (CirobuA:0000920). **Germ cells** (CirobuA:0000916) are maturing within the gonads.

The stalk base forms **test villi** (CirobuA:0000927), each one furnished with a **test vessel** (CirobuA:0000928) in continuity with the haemocele. They ensure firm adhesion to the substrate. In the body, several **blood sinuses** (CirobuA:0000856) can be recognized among organs.

After Stage 37, other territories become histologically recognizable (data not shown). These are the **cloacal cavity** (CirobuA:0000852), the **dorsal languets** (CirobuA:0000636) on the roof of the branchial chamber (CirobuA:0000863), the **pharyngo-epicardial openings** (CirobuA:0000868) putting the **epicardiac cavities** (CirobuA:0000879) in communication with the branchial one, the **endostylar appendix** (CirobuA:0000866), the **oral pigment spots** (CirobuA:0000899) and the **atrial pigment spots** (CirobuA:0000854) encircling the oral and the cloacal siphon border, respectively.

1.	Hirano, T. & Nishida, H. Developmental fates of larval tissues after metamorphosis in the ascidian, Halocynthia roretzi. II. Origin of endodermal tissues of the juvenile. *Dev. Genes Evol.* **210**, 55–63 (2000).

2.	Nakazawa, K. *et al.* Formation of the digestive tract in Ciona intestinalis includes two distinct morphogenic processes between its anterior and posterior parts. *Dev. Dyn.* **242**, 1172–83 (2013).

3.	Imai, J. H. & Meinertzhagen, I. A. Neurons of the Ascidian Larval Nervous System in Ciona intestinalis: I. Central Nervous System. *J. Comp. Neurol.* **501**, 316–334 (2007).

4.	Ryan, K., Lu, Z. & Meinertzhagen, I. A. The CNS connectome of a tadpole larva of Ciona intestinalis (L.) highlights sidedness in the brain of a chordate sibling. *Elife* **5**, 1–34 (2016).

5.	Moret, F. *et al.* The dopamine-synthesizing cells in the swimming larva of the tunicate Ciona intestinalis are located only in the hypothalamus-related domain of the sensory vesicle. *Eur. J. Neurosci.* **21**, 3043–3055 (2005).

6.	Razy-Krajka, F. *et al.* Monoaminergic modulation of photoreception in ascidian: evidence for a proto-hypothalamo-retinal territory. *BMC Biol* **10**, 45 (2012).

7.	Horie, T. *et al.* Regulatory cocktail for dopaminergic neurons in a protovertebrate identified by whole-embryo single-cell transcriptomics. *Genes Dev.* **32**, 1297–1302 (2018).

8.	Parrinello, D., Parisi, M., Parrinello, N. & Cammarata, M. Ciona robusta hemocyte populational dynamics and PO-dependent cytotoxic activity. *Dev. Comp. Immunol.* **103**, 103519 (2020).

9.	Bostwick, M. *et al.* Antagonistic Inhibitory Circuits Integrate Visual and Gravitactic Behaviors. *Curr. Biol.* 1–10 (2020). doi:10.1016/j.cub.2019.12.017

10.	Matsunobu, S. & Sasakura, Y. Time course for tail regression during metamorphosis of the ascidian Ciona intestinalis. *Dev. Biol.* **405**, 71–81 (2015).

11.	Cloney, R. A. Cytoplasmic filaments and morphogenesis: effects of cytochalasin B on contractile epidermal cells. *Zellforsch* **132**, 167–192 (1972).

12.	Numakunai, T. Hentai. in *Gendai Doubutsugaku No Kadai vol.5* (ed. Japan, T. Z. S. of) 135–175 (Gakkai Shuppan Center, 1977).

13.	Karaiskou, A., Swalla, B. J., Sasakura, Y. & Chambon, J.-P. P. Metamorphosis in solitary ascidians. *Genesis* **53**, 34–47 (2015).

14.	Lash, J. W., Cloney, R. A. & Minor, R. R. The effect of cytochalasin B upon tail resorption and metamorphosis in ten species of ascidians. *Biol. Bull.* **145**, 360–72 (1973).

15.	Terakubo, H. Q. *et al.* Network structure of projections extending from peripheral neurons in the tunic of ascidian larva. *Dev. Dyn.* **239**, 2278–87 (2010).

16.	Hirano, T. & Nishida, H. Developmental Fates of Larval Tissues after Metamorphosis in AscidianHalocynthia roretzi. *Dev. Biol.* **192**, 199–210 (1997).

17.	Chiba, S., Sasaki, A., Nakayama, A., Takamura, K. & Satoh, N. Development of Ciona intestinalis juveniles (through 2nd ascidian stage). *Zoolog. Sci.* **21**, 285–298 (2004).

18.	Davidson, B. Ciona intestinalis as a model for cardiac development. *Semin. Cell Dev. Biol.* **18**, 16–26 (2007).

19.	Davidson, B. & Levine, M. Evolutionary origins of the vertebrate heart: Specification of the cardiac lineage in Ciona intestinalis. *Proc. Natl. Acad. Sci. U. S. A.* **100**, 11469–73 (2003).

20.	Stolfi, A. *et al.* Early Chordate Origins of the Vertebrate Second Heart Field. *Science (80-. ).* **329**, 565 (2010).

21.	Wang, W., Razy-Krajka, F., Siu, E., Ketcham, A. & Christiaen, L. NK4 Antagonizes Tbx1/10 to Promote Cardiac versus Pharyngeal Muscle Fate in the Ascidian Second Heart Field. *PLoS Biol.* **11**, e1001725 (2013).

22.	Burighel, P., Cloney, R. A. & Cloney, B. Microscopic Anatomy of Invertebrates, Vol. 15. *Microsc. Anat. Invertebr.* **15**, 221–347 (1997).

23.	Kawamura, K. *et al.* Germline cell formation and gonad regeneration in solitary and colonial ascidians. *Dev. Dyn.* **240**, 299–308 (2011).

24.	Shirae-Kurabayashi, M. *et al.* Dynamic redistribution of vasa homolog and exclusion of somatic cell determinants during germ cell specification in Ciona intestinalis. *Development* **133**, 2683–93 (2006).

25.	Stolfi, A. *et al.* Divergent mechanisms regulate conserved cardiopharyngeal development and gene expression in distantly related ascidians. *Elife* **3**, e03728 (2014).

26.	Hozumi, A., Horie, T. & Sasakura, Y. Neuronal map reveals the highly regionalized pattern of the juvenile central nervous system of the ascidian Ciona intestinalis. *Dev. Dyn.* **244**, 1375–1393 (2015).

27.	Mackie, G. O., Burighel, P., Caicci, F. & Manni, L. Innervation of ascidian siphons and their responses to stimulation. *Can. J. Zool.* **84**, 1146–1162 (2006).

28.	Manni, L., Agnoletto, A., Zaniolo, G. & Burighel, P. Stomodeal and neurohypophysial placodes in Ciona Intestinalis: insights into the origin of the pituitary gland. *J. Exp. Zool. Part B Mol. Dev. Evol.* **304B**, 324–339 (2005).

29.	Veeman, M. T., Newman-Smith, E., El-Nachef, D. & Smith, W. C. The ascidian mouth opening is derived from the anterior neuropore: reassessing the mouth/neural tube relationship in chordate evolution. *Dev. Biol.* **344**, 138–49 (2010).

30.	Ogasawara, M. & Satoh, N. Isolation and Characterization of Endostyle-Specific Genes in the Ascidian Ciona intestinalis. *Biol. Bull.* **195**, 60–69 (1998).
