## Supplementary_File_03_Ciona_Staging_Table for "The ontology of the anatomy and development of the solitary ascidian *Ciona*"

**Table 1. Developmental Stages in *Ciona***
CirobuD:0000001

| Stages<br>St.1 - 26: Hotta et. al. 2007,<br>St. 26 - 37: present study | | Characteristics | Time after fertilization |
|---|---|---|---|

**Pre-embryonic development**
**CirobuD:0000002**

| | | |
|---|---|---|
| st.0 | Unfertilized egg | Spawned, not fertilized egg |

**Embryonic development, pre-metamorphosis**
**CirobuD:0000003**

| | | | |
|---|---|---|---|
| **I. Zygote period (0-1.0hr)** | | | |
| St. 1 | One cell | Zygote, fertilized egg | 24min (0.4hpf) |
| **II. Cleavage period (1.0-4.5hr)** | | | |
| St. 2 | 2-cell | Two cell-stage embryo | 55min (0.9hpf) |
| St. 3 | 4-cell | Four cell-stage embryo | 1hr 27min (1.45hpf) |
| St. 4 | 8-cell | Eight cell-stage embryo | 1hr 54min (1.9hpf) |
| St. 5a | early 16-cell | Early sixteen-cell stage embryo | 2hr 21min (2.35hpf) |
| St. 5b | late 16-cell | Late sixteen-cell stage embryo | 2hr 39min (2.65hpf) |
| St. 6a | early 32 cell | Early thirty two-cell stage embryo | 3hr (3hpf) |
| St. 6b | late 32 cell | Late thirty two-cell stage embryo | 3hr 12min (3.2hpf) |
| St. 7 | 44-cell | Fourty four-cell stage embryo. The vegetal side of the embryo is round | 3hr 21min (3.35hpf) |
| St. 8 | 64-cell | Sixty four-cell stage embryo. Embryo has a square shape when seen form the top, with bulging B7.4 cells | 4hr (4hpf) |
| St. 9 | 76-cell | Seventy six cell stage embryo. The vegetal side of the embryo is flat | 4hr 12min (4.2hpf) |
| **III. Gastrula Period (4.5-6.3hr)** | | | |
| St. 10 | 110-cell, initial gastrula | Gastrulation starts with the apical constriction of A7.1 blastomeres | 4hr 33min (4.5hpf) |
| St. 11 | early gastrula | The ntochord has invaginated. The vegetal side of the embryo has a horseshoe shape. | 4hr 54min (4.9hpf) |
| St. 12 | mid gastrula | Six-row neural plate stage. The blastopore is still central and open | 5hr 39min (5.65hpf) |
| St. 13 | late gastrula | The blastopore is in posterior position and nearly closed. The embryo elongates anteriorly. The neural plate has more than 6 rows and the A-line neural rows (I and II) start to curve (beginning of neurulation). The large b6.5 progeny are coming together at the midline | 5hr 55min (5.9hpf) |
| **IV. Neurula Period (6.3-8.5hr)** | | | |
| St. 14 | early neurula | A-line neural plate forms a groove lined by b6.5 descendants. The embryo has a diamond shape. The groove is not | 6hr 21min (6.35hpf) |
| St. 15 | mid neurula | The neural tube has formed on most of its length. The embryo has an oval shape. The a-line neural plate also forms a | 6hr 48min (6.8hpf) |
| St. 16 | late neurula | The neural tube starts to form in the posterior territories. The embryo elongates | 7hr 24min (7.4hpf) |
| **V. Tailbud Period (8.5-17.5hr)** | | | |
| St. 17 | initial tailbud I | First indication of a separation between tail and trunk territories. The tail is not bent and has the same length as the trunk. Intercalation of notochord cells is not concluded | 8hr 27min (8.45hpf) |
| St. 18 | initial tailbud II | The tail is clearly separated from the trunk. Tail and trunk have same length. Neuropore still open, a-line neurulation | 8hr 50min (8.8hpf) |
| St. 19 | early tailbud I | The tail is angled about 40°and is slightly longer than the trunk. A few anterior most notochord cells begin to intercalate and arrange in line | 9hr 19min (9.3hpf) |
| St. 20 | early tailbud II | Neuropore closed, tail angled about 60°, neurulation complete | 9hr 30min (9.5hpf) |
| St. 21 | mid tailbud I | Tail 1 1/2 times longer than trunk and curve ventrally (90°). Intercalation of notochord cells just finished | 10hr 2min (10hpf) |
| St. 22 | mid tailbud II | The body adopts a half circle shape. Tail twice as long as trunk. | 10hr 54min (10.9hpf) |
| St. 23 | late tailbud I | Initiation of the pigmentation of the otolith. Tail strongly curved with tip close to the anterior end of the trunk | 11hr 54min (11.9hpf) |
| St. 24 | late tailbud II | Notochord vacuolation begins, palps start to be visible at the front end of the embryo. Tail straightens | 13hr 27min (13.5hpf) |
| St. 25 | late tailbud III | Ocellus melanization. All notochord cells have vacuoles. Tail bent dorsally | 15hr 54min (15.9hpf) |

| | | | | | |
|---|---|---|---|---|---|
| **VI. Larva Period (St.26~29, 17.5~24hpf)*** | | | | **Chiba's Stage (2004)** | ANISEED (2017) |
| St. 26<br>CirobuD:0000049 | hatching larva | Hatching, spherical head shape, immature papillae with pyramidal shape, irregular tail movements | 17hr 30min (17.5hpf) | Stage0 | St. 26 |
| St. 27<br>CirobuD:0000050 | early swimming larva | Spindle-like head shape, regular tail movements and swimming behaviour | 17.5-20 hpf | | St. 27 |
| St. 28<br>CirobuD:0000051 | mid swimming larava | Elongated papillae and expansion of their basal part, square head, spherical test cells, cilia in epidermal sensory neurons recognizable, preoral lobe recognizable | 20-22hpf | Stage1 | St. 28 |
| St. 29<br>CirobuD:0000052 | late swimming larva | Longer and narrower head with respect to St. 28, trunk profile squared at transition between trunk and tail | 22-24hpf | Stage2 | St. 29, 30 |

**Metamorphosis**
**CirobuD:0000004**

| | | | | | |
|---|---|---|---|---|---|
| **VII. Adhesion period (St.30, 24~27hpf)** | | | | | |
| St. 30<br>CirobuD:0000053 | adhesion | Curved papillae, otolith and ocellus remnants recognizable | 24-27 hpf | | St. 31, 32 |
| **VIII.Tail absorption period (St.31~33, 27~30hpf)** | | | | | |
| St. 31<br>CirobuD:0000054 | early tail absorption | Beginnng of tail absorption, tail bending at the transition between trunk and tail, otolith and ocellus remnants recognizable | 27 hpf | | St. 33 |
| St. 32<br>CirobuD:0000055 | mid tail absorption | 50% of tail absorbed into trunk. Tail shrinked and thickened, otolith and ocellus remnants recognizable | 28 hpf | | St. 34 |
| St. 33<br>CirobuD:0000056 | late tail absorption | Tail completely absorbed, papiliae no more recognizable, otolith and ocellus remnants recognizable | 29 hpf | | St. 35 |
| **IX. Body axis rotation period (St.34~36, 30~60hpf)** | | | | | |
| St. 34<br>CirobuD:0000057 | early body axis rotation | Beginning of body axis rotation (angle between the stalk and the endostyle more than 0°), outer tunic compartment and outer cuticle layer no more present, tunic cells recognizable in definitive tunic, otolith and ocellus remnants recognizable | 30-36 hpf | Stage3a | St. 36 |
| St. 35<br>CirobuD:0000058 | mid body axis rotation | Body axis rotation of 30°- 60°, one pair of gill-slit recognizable, otolith and ocellus remnants recognizable | 36-45 hpf | | St. 37, 38 |
| St. 36<br>CirobuD:0000059 | late body axis rotation | Two pair of gill-slit open, body axis rotation at 80°-90°, filtering and feeding activity present, otolith and ocellus remnants recognizable, heart beating | 45-60 hpf | Stage3b | St. 39, 40 |

**Post-metamorphosis**
**CirobuD:0000005**

| | | | | | |
|---|---|---|---|---|---|
| **X. Juvenile period (St.37~41, 60hpf~)** | | | | | |
| St. 37<br>CirobuD:0000060 | early juvenile I | Body axis rotation completed, stomach swollen, otolith and ocellus remnants recognizable | 63-72 hpf (3dpf) | Stage4 | St. 41, 42 |
| St. 38<br>CirobuD:0000061 | early juvenile II | Larval tail remnants totally adsorbed | 3-4 dpf | Stage5 | St. 43, 44 |
| St. 39<br>CirobuD:0000062 | mid juvenile I | Additional gill slit begin to open, appearance of stomach, gut and neural grand | 4-6 dpf | | St. 45 |
| St.40<br>CirobuD:0000063 | mid juvenile II | Gonad in form of oval vesicle (corresponding to Stage 6 in Chiba et. al., 2004) | 6-7dpf | Stage6 | St. 46 |
| St.41<br>CirobuD:0000064 | late juvenile | Atrial siphon begins to fuse (corresponding to Stage 7 in Chiba et. al., 2004) | 7dpf~ | Stage7 | St. 47 |
| **XI. Young adult period** | | | | | |
| St.42 | | 2nd Ascidian Stage (corresponding to Stage 8 in Chiba et. al., 2004) | | Stage8 | |
| **XII. Mature adult period** | | | | | |
| St.43 | | Gonad fully matured | | Adult | |

* The duration of larval swimming differs among individuals. Matsunobu et al. (2015) showed that the hatched larva requires at least three or four hours to get competence to commence metamorphosis. So the time after fertilization during Larva Period was broad.
